# Conserved patterns of mammal–microbe codiversification and interspecific bacterial sharing

**DOI:** 10.64898/2026.10.16.759742

**Authors:** Gabriele Schiro, Aishwarya Patel, Ellen Tierney, Bret Pasch, Nathan S. Upham, Taichi A. Suzuki

**Affiliations:** Biodesign Center for Health Through Microbiomes, Arizona State University, Tempe, AZ, USA; Department of Ecology and Evolutionary Biology, University of Arizona, Tucson, AZ, USA; School of Natural Resources and the Environment, The University of Arizona, Tucson, AZ, USA; School of Life Sciences, Arizona State University, Tempe, AZ, USA; College of Health Solutions, Arizona State University, Tempe, AZ, USA

## Abstract

What determines whether a gut microbial lineage has fidelity to one host species or is shared freely across multiple species? Factors that predict microbial community composition, including host species, diet, and environmental variation, have been well documented in wild animals. However, the eco-evolutionary processes maintaining host species- or population-specific microbial lineages remain unresolved. Here, we analyze 519 gut microbiome samples collected from 15 species of wild rodents across two mountains in the Madrean Sky Islands of southeastern Arizona using full-length 16S rRNA sequencing. This montane system provides a natural experiment in which adjacent replicated elevational gradients bring different combinations of rodent species into sympatry, allowing host evolutionary history to be disentangled from shared environmental effects. We find that host phylogeny, rather than environment, is the primary driver of microbial strain sharing across host species. We recover replicated signatures of codiversification on both mountains, indicating shared evolutionary history between hosts and their microbes, despite ample opportunities for cross-species environmental transmission. This evolutionary coupling is associated with predicted differences in bacterial oxygen tolerance and optimal growth temperature: codiversifying lineages are predominantly obligate anaerobes with optimal growth near host body temperature, suggesting more limited out-of-host persistence than non-codiversifying taxa. This pattern mirrors findings from the human gut microbiome, suggesting conserved constraints on microbial dispersal across mammalian hosts. The cost-effectiveness of long-read amplicon sequencing for phylogenetic inference within bacterial species opens new opportunities for large-scale evolutionary and ecological studies of host-microbe interactions in wildlife.

## Introduction

The mammalian gut microbiome exhibits variation across multiple scales of organization, ranging from differences among individuals within a species to divergence across entire orders (Ley et al., 2008; Ma, 2021), with profound implications for host biology (McFall-Ngai et al., 2013; Suzuki, 2017). At the community level within hosts, the factors structuring these patterns have been extensively studied (Kartzinel et al., 2019; Kohl, 2020; Levin et al., 2021; Wu et al., 2022), but variation at the strain level remains comparatively underexplored. Microbial strains within the same species can differ markedly in their functional diversity (Van Rossum et al., 2020), and accumulating evidence suggests that host populations and species harbor distinct strain-level variation shaped by their evolutionary histories (Andreu-Sánchez et al., 2025; Moeller et al., 2013, 2017). Yet, how these strains are transmitted across hosts, particularly over deeper evolutionary timescales and across ecologically diverse settings, remains poorly understood.

Microbial transmission is fundamental to shaping microbiome assembly and evolution, as gut microbes must first be acquired before they can persist within hosts (Moeller et al., 2018). To date, research has largely focused on elucidating microbial transmission routes within populations of the same species, including humans and wild animals (Labisa-Morais et al., 2025; Petrullo et al., 2025; Raulo et al., 2024; Valles-Colomer et al., 2023). Although strain sharing across species has been investigated (Dillard et al., 2026; Moeller et al., 2013, 2017), the determinants of microbial transmission across species remain relatively underexplored. Microbes that are consistently transmitted within host populations but not between host species are more likely to form evolutionary stable associations with their hosts, creating greater opportunities for host-microbial coevolution (Groussin et al., 2020; Sanders et al., 2023; Suzuki et al., 2022). Previous studies in humans (Suzuki et al., 2022), primates (Sanders et al., 2023), and rodents (Sprockett et al., 2025) have revealed patterns of host–microbial codiversification across specific microbial clades, where microbial lineages have diversified in parallel with their hosts, a signal reflected by congruent host and microbial phylogenies (Davenport et al., 2017). Understanding what maintains host-specific microbial lineages across evolutionary time has broad implications for animal ecology and evolution, from disease dynamics to microbiome-based interventions (Groussin et al., 2020).

Microbial traits are likely a key, yet underappreciated, determinant of these transmission and persistence patterns. Aerotolerant and spore-forming taxa are better equipped to persist in external environments, potentially facilitating transmission across individuals and species (Dillard et al., 2026; Raulo et al., 2024). Within populations, aerotolerant taxa are more frequently associated with indirect and social transmission in rodents (Petrullo et al., 2025; Raulo et al., 2024). In contrast, strictly anaerobic and non-spore-forming microbes may rely more heavily on direct host-to-host transmission, which is more frequent among kin and conspecifics (Raulo et al., 2024; Valles-Colomer et al., 2023), potentially contributing to the host–microbial codiversification patterns observed in mammals (Suzuki et al., 2022). Consistent with this, bacteria that exhibit long-term codiversification with their hosts have been associated with the loss of traits such as aerotolerance and temperature tolerance, reflecting adaptation to the gut environment and reduced capacity for out-of-host persistence (Moeller et al., 2016; Suzuki et al., 2022). Whether these microbial trait-transmission relationships extend across host species boundaries remains largely unexplored.

Despite the emerging links between microbial trait evolution and transmission mode, the processes that generate and maintain host species- or population-specific microbiomes, particularly at the strain level, remain unresolved. Closely related hosts often share diets, habitats, and environmental microbial sources, complicating efforts to determine whether microbial lineage sharing across hosts arises from shared ecology, host evolutionary history, or ongoing microbial exchange. This problem is especially pronounced in most existing studies that compare hosts in allopatry, limiting the ability to distinguish between transmission within host lineages (e.g., vertical or familial transmission) and acquisition from shared environments. Notably, primates and rodents in captivity harbor different microbial communities and strains than their wild counterparts (Moeller, 2025; Nishida & Ochman, 2021), highlighting the importance of studying sympatric host populations in natural settings. Yet even in field-based studies, observations are inherently noisy and often lack replication, emphasizing the need for natural experimental systems with independent biological replicates. Finally, commonly used short-read 16S rRNA sequencing lacks the resolution to resolve subspecies or strain-level variation (Johnson et al., 2019), the scale at which transmission dynamics and codiversification signals can be resolved. Although shotgun metagenomics provides the necessary resolution, its high cost limits application to large sample sizes, a critical constraint when studying diverse wildlife communities.

In this study, we leverage an unparalleled natural experimental design, drawing on a dataset of > 500 gut microbiomes from 15 rodent species across two Madrean Sky Islands in southeastern Arizona (USA) (**Fig. 1A-C**). Across both mountain ranges, multiple host species co-occur in sympatry along shared elevational gradients, enabling replicated comparisons of closely and distantly related species across similar and different vegetation zones (**Fig. 1D-E**). This design allows us to simultaneously disentangle the effects of host phylogenetic relatedness, shared vegetation type, and geographic separation across two independent locations. These high-elevation forests are separated by desert lowlands that have maintained population isolation for approximately 10,000–50,000 years (Betancourt et al., 1990), a timescale comparable to human population divergence times, making them ideal for testing the mechanisms driving host– microbial associations. Using full-length 16S rRNA sequencing, we address three main questions: (1) What are the primary determinants of microbial strain sharing across host species including host phylogeny, vegetation similarity, and geography? (2) Do replicated signatures of host– microbial codiversification emerge across two mountain ranges independently? And (3) which microbial traits associated with environmental persistence and transmission underlie these patterns?

**Figure 1.**
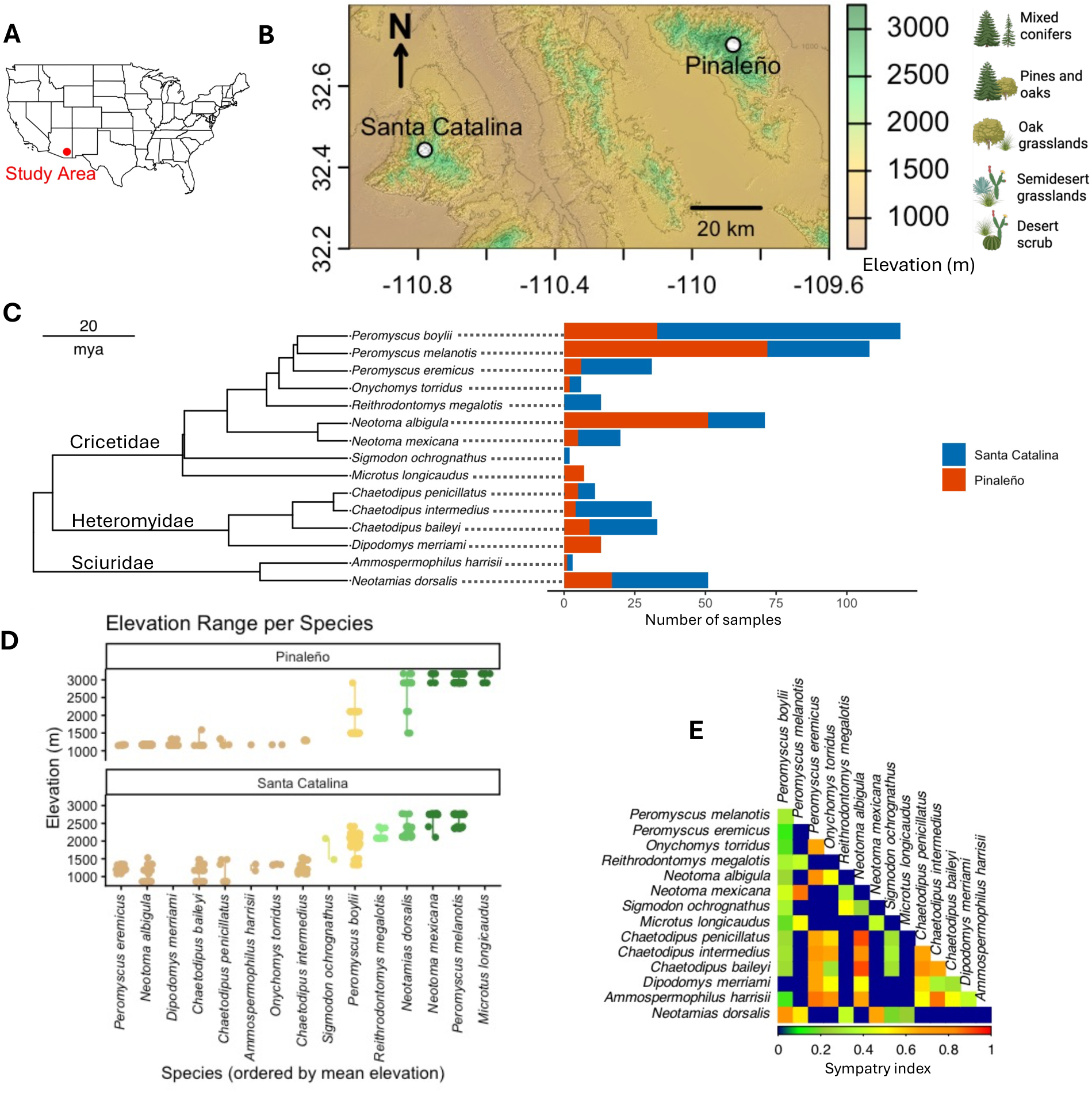
Rodent communities across sky islands exhibit diverse combinations of phylogenetic relatedness, geographic overlap, and environmental isolation by elevation. (A) Location of the study area in the southwestern United States. (B) Topographic map of the Santa Catalina and Pinaleño Mountains showing elevation gradients and indicative vegetation encountered. (C) Phylogeny of sampled species (left) and sample sizes by mountain range (right; blue = Santa Catalina, red = Pinaleño). (D) Elevational distributions of species in the Pinaleño (top) and Santa Catalina (bottom) Mountains, with species ordered by mean elevation and color scaled as in panel B. (E) Heatmap of pairwise co-occurrence (similarity) among species, calculated as the proportion of sampling locations where both species were trapped.

## Materials and Methods

### Sample collection

Fecal and tissue samples from small mammals were collected across multiple sites in both the Pinaleño and Santa Catalina mountains (**Fig. 1A-B**) between November 2021 and June 2025. In total, 524 samples from 15 species across three rodent families were collected from two mountain ranges (**Fig. 1C**), capturing both conspecifics occupying distinct vegetation zones and heterospecifics co-occurring within the same zones, allowing us to disentangle the determinants of strain sharing (**Fig. 1D**). For example, *Peromyscus* species show a stratified distribution: *P. eremicus* at low elevations, *P. boylii* at mid elevations, and *P. melanotis* at high elevations, with P. boylii overlapping in habitat with both other species. Together, this sampling design spans a wide range of phylogenetic relatedness, geographic isolation, and vegetation similarity (**Fig. 1E**).

To minimize the effects of seasonal variation, animals were collected multiple times from the same location across seasons and years. We used Sherman live traps baited with sterilized birdseed, oats, and peanut butter to collect animals overnight. Microhabitats included desert, grassland, chaparral, oak forest, and conifer forest. Upon capture, we measured body weight, tail length, and hind foot length and collected fecal pellets and an ear tissue biopsy. Animals were then released at their capture sites or euthanized. Fecal samples were collected within 18 hours of capture and immediately stored on dry ice or in liquid nitrogen until transported to the University of Arizona or Arizona State University and stored at −80°C until further processing. Detailed metadata for each sample are listed in **supplementary table S1**. All procedures involving animals were approved under the Arizona Game and Fish Department collection permits (SP163376, SP407147, SP623885, SP939945, SP483771, SP055536), the National Forest Service permits (SUP2557, SAN2139, SAN2218, SAN2329, Mount Graham biology program), and by the Institutional Animal Care and Use Committees at the University of Arizona (2022-0864) and Arizona State University (25-2163R, 21-1849R).

### DNA extraction and sequencing

DNA was extracted from fecal samples using the DNeasy PowerSoil Kit (QIAGEN) following the manufacturer’s instructions. 16S Kinnex libraries were generated following the manufacturer’s protocol at the Genomics Core at Arizona State University. DNA samples were diluted to less than 20 ng/uL and the full-length 16S rRNA gene was amplified from microbial DNA with the barcoded 27f/1492r Kinnex primer set. Amplicons were pooled in equal volumes and a SMRTbell cleanup was performed. Kinnex adapters were added via PCR, followed by pooling and an additional SMRTbell cleanup. Kinnex Arrays were then assembled to concatenate the 16S amplicons into a linear array, followed by a SMRTbell cleanup and nuclease treatment. The final SMRTbell library was cleaned up using 3.1X AmPure® PB beads, quantified with a Qubit fluorometer, and assessed for quality using a Femto Pulse system prior to equimolar pooling of barcoded libraries. Pooled libraries were sequenced on a PacBio Revio SMRT cell. All samples were randomized prior to library preparation. Twenty negative controls were included across sequencing runs and returned no visible PCR product and no reads. To confirm species identities of *Peromyscus*, two samples from a single sampling site were randomly selected for *cytochrome b* gene amplification using the primers described by Naidu et al. (2012). Amplicons were sequenced at the University of Arizona Genetics Core. Resulting sequences were compared against the NCBI GenBank database using the Basic Local Alignment Search Tool (BLAST; blastn), and species identities were confirmed based on the top hit.

### ASV inference

Amplicon sequence variants (ASVs) were inferred using the HiFi-16S-workflow (pb-16-nf V0.9 https://github.com/PacificBiosciences/HiFi-16S-workflow), with the following modifications. The DADA2 maximum expected error rate was set to four instead of the default two, to account for the longer read length. Taxonomy was assigned with the RDP naive Bayes classifier, trained on the GTDB database release 226 (Parks et al., 2022). Samples with fewer than 10,000 reads (n = 5) were excluded from downstream analysis.

### Host phylogenetics and dietary traits

All analyses were conducted in R using the *ape* package (Paradis et al., 2004). We obtained a posterior sample of 1,000 phylogenetic trees for the rodent species in the study from vertlife [https://vertlife.org/phylosubsets/ (Upham et al., 2019)]. A majority-rule consensus tree was constructed from the tree sample using the “consensus” function. Consensus branch structure and edge relationships were further calculated using “consensus.edges”. Node support values were calculated by estimating clade frequencies across the full set of trees using the “prop.clades” function, which assigns to each node the proportion of trees in which the corresponding clade is recovered. All nodes in the resulting consensus tree showed maximal support (100%). Dietary traits for each species in the database were retrieved from the Elton traits database (Wilman et al., 2014).

### Microbial phylogenetics

To construct a phylogenetic tree of ASVs, we first removed ASVs with very low abundance and prevalence from the dataset (retaining only those with at least 50 counts across all samples and a minimum abundance of at least 10 reads in at least one sample). This step reduced computational time and provided a way to control for low-abundance sequencing errors not fully corrected during the DADA2 denoising step, without affecting alpha and beta diversity patterns (**supplementary figure S1**). The remaining 47,790 ASVs were aligned using the tool SINA (Pruesse et al., 2012), which uses multiple reference sequences from the SILVA v 138.2 SSURef NR99 database (Quast et al., 2012) as a reference for alignment of candidate sequences, using default parameters. Multiple sequence alignments were trimmed using trimAl (Capella-Gutiérrez et al., 2009) to remove poorly aligned regions. Alignment columns with more than 80% gap occupancy were discarded, and the remaining alignment was input into FastTree under the GTR+CAT model with the -fastest flag disabled to improve topological accuracy.

### Alpha and beta diversity

Alpha and beta diversity were calculated on rarefied reads (10,000) using the filtered ASV table. Throughout this manuscript, we use “strain” operationally to refer to bacterial sequence variants distinguished by full-length 16S rRNA ASVs, which serve as proxies for within-species variation. ASVs do not necessarily correspond one-to-one with genome-defined strains because individual genomes can contain multiple 16S rRNA variants and distinct strains can share identical 16S rRNA sequences. To investigate the drivers of bacterial strain sharing across Sky Islands rodents, we computed pairwise Sørensen similarity for all individual pairs, using shared ASVs as a proxy for strain sharing (n = 115,963 pairs). To test for the effect of each predictor on microbial community dissimilarities (Sørensen-Dice and Bray-Curtis), we ran a PERMANOVA [function adonis2 in vegan (Oksanen et al., 2018)]. Predictors included host species identity, mountain range, elevation, and vegetation greenness (NDVI). To isolate the effect of vegetation independent of elevation, we calculated the residuals of a linear model regressing mean NDVI (averaged across 2022–2025 Sentinel-2 imagery collected in June) on elevation and used these residuals as the NDVI predictor in all models.

### Annotation of microbial traits

To infer microbial traits, we used the trait dataset from metaTraits (Podlesny et al., 2026). Briefly, family-, genus-, and species-level trait summary tables were downloaded and traits of interest were extracted. We selected three binary traits (obligate anaerobiosis, Gram stain, and sporulation) and three continuous traits (genome size, GC content and optimal growth temperature), based on their relevance to out-of-host environmental persistence, previous evidence linking them to host–microbe codiversification, and their broad representation in the database as traits routinely recorded during isolate characterization (McCutcheon & Moran, 2012; McDADE & Hall, 1964; Suzuki et al., 2022; Teng et al., 2023). For binary traits, we extracted data from the database and calculated the percentage of reported strains within each taxonomic group displaying that trait. We used this percentage as the estimated probability of an ASV or taxonomic group exhibiting the trait (e.g., the percentage of strains classified as strictly anaerobic within a family). For continuous traits, the median value across strains within each taxonomic group was extracted. Trait values were also assigned to ASVs hierarchically, first at the species level, and then at the genus level, and then at the family level when finer-level trait data were unavailable or the ASV lacked a finer-level taxonomic assignment.

### Modelling of host species occupancy as a function of microbial traits

We modeled ASV host occupancy using a binomial generalized linear model, with the response expressed as the number of host species in which each ASV was detected relative to the total number of host species sampled. Categorical predictors (obligate anaerobiosis, sporulation, and Gram stain) were treated as proportional variables scaled between 0 and 1 using min–max normalization to facilitate comparison of effect sizes across traits. Continuous predictors (genome size, optimal growth temperature (OGT), and GC content) were standardized to unit variance, and quadratic terms were included to account for potential nonlinear relationships with occupancy. Multicollinearity among predictors was assessed using variance inflation factors. An equivalent model fitted using raw, unscaled trait values was used for plotting predicted relationships. To account for shared evolutionary history, we additionally fitted an equivalent phylogenetic generalized linear mixed model (PGLMM) using the phyr package (Li et al., 2020). Owing to computational constraints, this analysis was restricted to the 5,000 most abundant ASVs.

### Bayesian multi-membership regression model

To test the effect of each predictor on ASV sharing rate, we fitted an ordered beta regression model using the ordbetareg package (Kubinec, 2023), an extension of Bayesian multimembership regression models (Bürkner, 2017) designed for continuous response variables bounded between 0 and 1. Sørensen-Dice was modeled as a function of the following fixed effects: phylogenetic distance, elevational distance (as a proxy for vegetation zone), NDVI residualized on elevation (to capture vegetation productivity independent of elevation zone), and geographic distance between host pairs. To account for non-independence in the data, we included multi-membership random intercepts for paired samples and paired species (Raulo et al., 2024). The model was estimated in Stan (Raulo et al., 2024) via Hamiltonian Monte Carlo with 4 chains, 3,000 iterations per chain, and 1,000 warmup iterations, using 4 cores and within-chain threading across 4 threads. Same-species comparisons were excluded before fitting the model.

### Codiversification test

We investigated patterns of codiversification between host and bacterial lineages by testing phylogenetic nodes across the distal quarter of the bacterial phylogeny, following Sprockett et al. (2025). To ensure that analyses focused on well-represented bacterial clades rather than nodes dominated by a single highly abundant ASV, we restricted analyses to nodes containing at least five distinct dominant ASVs, where each dominant ASV represented the highest relative-abundance ASV in at least one sample within that node. In addition, nodes were required to be present in at least 50 samples and across a minimum of five host species, leaving a total of 4,467 nodes to be tested. For each sample, the ASV within that node with the highest relative abundance was considered representative for that sample. Spearman and Pearson Mantel tests were performed for each of the 4,467 filtered nodes, with Spearman used for downstream analysis, as it is less sensitive to outliers and does not assume linearity. P-values were corrected using the Benjamini-Hochberg false discovery rate method. To reduce redundancy among overlapping nodes, nodes sharing at least 75% of their tips were identified, and only the node with the stronger Mantel correlation was retained. Only nodes with Mantel rho >0.75 and an adjusted p-value < 0.05 were considered strongly codiversifying. For comparison, nodes with rho < 0.25 and no p-value threshold were classified as lacking evidence for codiversification, as this excludes weak but statistically significant correlations and provides a more conservative comparison group. Mantel tests were also performed separately for each mountain range to verify reproducibility of codiversification patterns across the two geographically separated locations.

### Association of microbial traits with codiversification

To test whether codiversification was associated with specific bacterial traits, we grouped ASVs by family and, for each family represented by at least 10 ASVs, calculated the number of ASVs falling within strongly codiversifying nodes. We then calculated an observed test statistic that was compared with a null distribution generated by permutation. For binary traits (obligate anaerobiosis, sporulation, and Gram stain) and GC content, the observed statistic was the weighted mean of the family-level trait values, with each family weighted by its number of codiversifying ASVs. For optimal growth temperature, the observed statistic was the weighted mean absolute deviation from host body temperature (37 °C). To check for possible phylogenetic conservatism of traits between families, we used the D statistic for binary traits (obligate anaerobiosis, sporulation), and Pagel’s lambda for the continuous trait of mean optimal growth temperature, with significance assessed via 1,000 permutations (D statistic) or a likelihood-ratio test (lambda).

Statistical significance was assessed using 10,000 permutations, randomly re-assigning the association between family-level trait values and codiversification labels to generate a null distribution for each test statistic. P-values were calculated as the proportion of permutations yielding a statistic equal to or more extreme than the observed value. One-sided tests were used for the enrichment of obligate anaerobes, reduced deviation from host body temperature, and reduced genome-size variation, whereas two-sided tests were used for GC content, sporulation, and Gram-positive proportion, for which no directional hypotheses were specified.

## Results

### Host species identity is the dominant predictor of gut microbiome composition

Across 519 samples from 15 rodent species across two mountains in our dataset (**Fig. 1**), bacterial taxa belonged to 47,790 different amplicon sequence variants (ASVs). At the phylum level, most ASVs belonged to Bacillota_A (35.5% relative abundance), Bacteroidota (30.9%) and Bacillota (27.3%). At the family level, the most abundant families were Muribaculaceae (23.5%), followed by Lactobacillaceae (21.8%) and Lachnospiraceae (19.3%). Regardless of the mountain sampled, Heteromyidae (genera *Chaetodipus* and *Dipodomys*) were consistently depleted of Lactobacillaceae relative to other samples (**supplementary figure S2**). In a model accounting for mountain range, elevation, and vegetation greenness, species identity was the strongest predictor of gut microbiome richness (F₁₄,₅₀₁ = 36.61, p < 0.001, **supplementary figure S3**), followed by mountain range (F₁,₅₀₁ = 4.09, p = 0.044), while elevation and vegetation greenness did not significantly contribute (elevation: F₁,₅₀₁ = 0.03, p = 0.867; NDVI: F₁,₅₀₁ = 0.46, p = 0.497). When species was not included as a predictor, microbiome richness differed significantly by mountain range and showed a nonlinear relationship with elevation, peaking at mid-elevations (linear term: β = 0.393, p < 0.001, AIC = 6968.37; quadratic term: β = −0.0000759, p = 0.003, AIC = 6961.74; **Fig. 2A**). Richness was lower in the Santa Catalina Mountains (β = −79.20, p < 0.001) while vegetation greenness did not significantly predict richness (β = 47.0, p = 0.627), and the quadratic elevation terms indicated a hump-shaped relationship, with richness increasing at lower elevations and declining at higher elevations (linear term: β = 0.393, p < 0.001; quadratic term: β = −0.0000759, p = 0.003).

**Figure 2.**
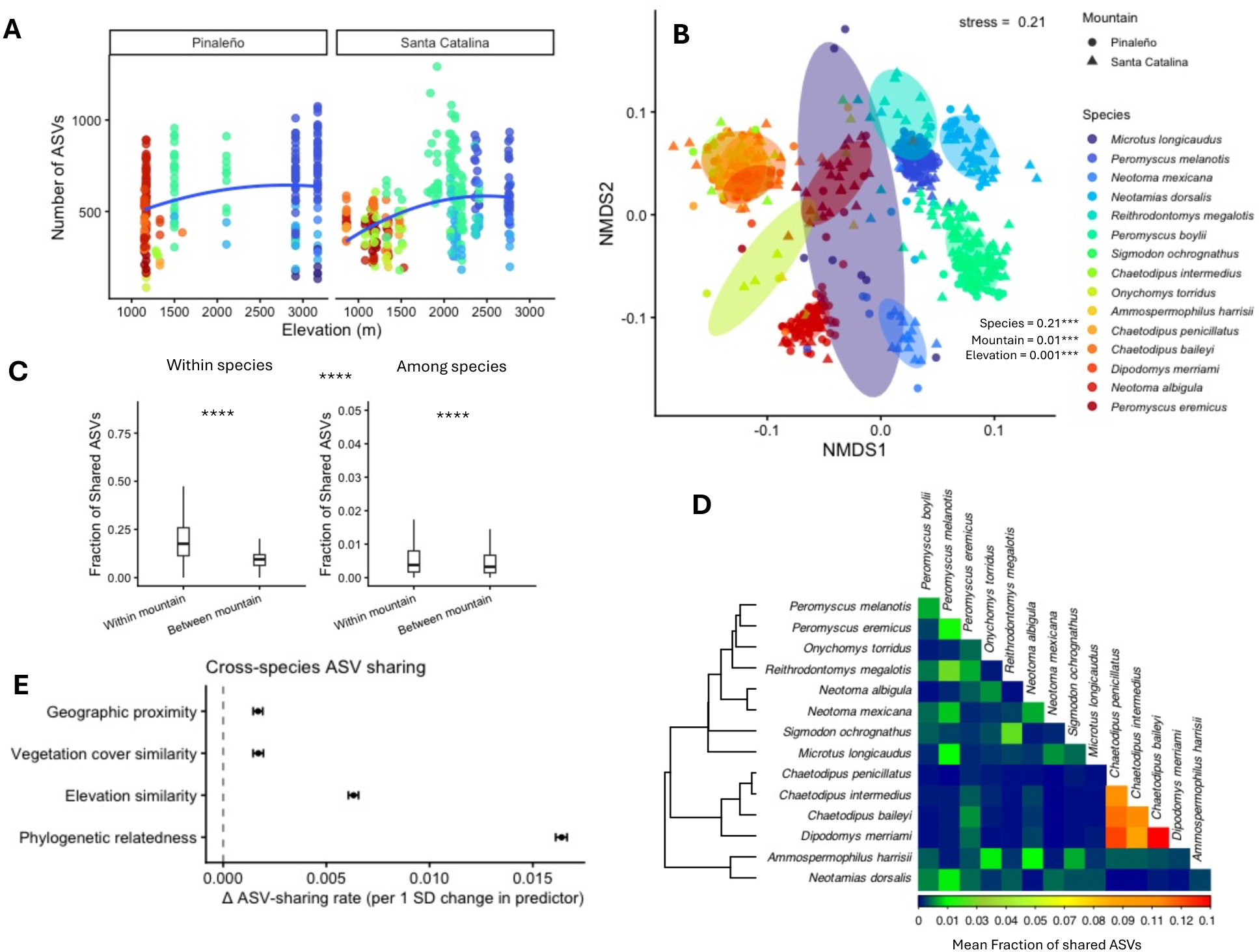
Species and elevation are stronger predictors of community composition than mountain. (A) Relationship between elevation (meters above sea level) and microbial richness (number of ASVs) for each mountain range, with fitted trend lines illustrating diversity patterns along elevation gradients. (B) Non-metric multidimensional scaling (NMDS) ordination of presence-absence microbial community similarities (Sørensen-Dice index), showing clustering by host species and differences between mountain ranges (circles = Pinaleño; triangles = Santa Catalina). PERMANOVA pseudo-R squared values and p-values are also reported (*** P < 0.001). (C) Distribution of pairwise Sørensen similarity (proportion of shared ASVs) between individual rodents, faceted by comparisons within and between mountain ranges for both within-species comparisons (left panel) and among-species comparisons (right panel; note different Y-axis scales). P-values are calculated using a permutation test in which species and mountain labels are randomly shuffled among samples and the recalculated mean is compared with the observed one (9999 permutations, **** P < 0.0001). (D) Mean pairwise fraction of shared ASVs between all species pairs (n = 15 species), ordered by host phylogeny. Warmer colors indicate a higher rate of microbial strain sharing. (E) Results of Bayesian multi-membership regression models estimating the effect of phylogenetic relatedness, elevational similarity, vegetation cover similarity (residuals of NDVI regressed on elevation) and geographic proximity on pairwise ASV sharing across host species pairs.

To capture strain sharing patterns, we used Sørensen-Dice similarity, a presence– absence-based metric particularly well suited to full-length 16S rRNA ASV data. Consistent with the microbial richness results, we found that species was the strongest determinant of taxon sharing among samples (**Fig. 2B**). PERMANOVA (adonis2; 999 permutations, each term tested while controlling for all others) showed significant effects of species (R² = 0.207, F = 10.43, p = 0.001), mountain (R² = 0.012, F = 8.62, p = 0.001), elevation (R² = 0.005, F = 3.31, p = 0.001), and vegetation greenness independent of elevation (R² = 0.007, F = 5.23, p = 0.001), indicating that strain sharing was strongly structured by host species identity: individuals of the same species shared a substantially higher fraction of ASVs (permutation Wilcoxon test, *Z* = 285.47, p < 0.0001, **Fig. 2C**). Among same-species pairs, individuals from the same mountain shared more ASVs than those from different mountains, and this geographic signal was also evident in cross-species comparisons (**Fig. 2C**). When strain sharing was restricted to individual species, mountain range had a significant effect in all tested species except *C. penicillatus* (PERMANOVA, p < 0.05), with effect sizes ranging from R² = 0.05 to 0.26 (**supplementary table S2**). Species at higher elevations tended to show stronger geographic structuring, although the trend was not significant (r = 0.45, p = 0.25, **supplementary figure S4**). Overall, these results indicate that geographic structuring of microbial communities is evident within host species but comparatively weak across species, where host identity is the dominant driver. Similar but weaker patterns were observed for Bray–Curtis dissimilarities (**supplementary figure S5**), indicating that although similar taxa are shared among samples, their relative abundances vary, suggesting that abundance patterns may be more strongly influenced by ecological factors not captured in our experimental design, such as seasonality and interannual variations.

### Phylogenetic relatedness is the strongest determinant of ASV sharing across species

Having established that host species identity structures both microbial richness and community composition, we next asked what drives strain sharing specifically across host species. To this end, we calculated the proportion of ASVs shared between species pairs (**Fig. 2D**), and modeled this using a Bayesian multi-membership regression framework that explicitly includes phylogenetic relatedness as a predictor, moving beyond species identity to quantify the continuous effect of host evolutionary history on strain sharing. Phylogenetically similar host species shared a higher fraction of ASVs, consistent with phylosymbiosis (**supplementary figure S6**). Notably, Heteromyidae appeared to behave differently from other groups, sharing a higher proportion of ASVs among themselves, suggesting an additional role of geographic proximity beyond host identity alone. This approach accounts for pairwise distances while controlling for the non-independence of samples and species pairs, and same-species pairs were excluded to focus on interspecific transmission and avoid inflating the phylogenetic signal. Host phylogeny was the strongest predictor of ASV sharing, followed by vegetation zone (elevational distance and NDVI residuals) and geographic proximity (**Fig. 2E**, with **supplementary table S3** providing the full model output). In other words, at a standardized unit of variation for each predictor, more closely related host species tended to share a greater fraction of ASVs than species that diverged more deeply in evolutionary time. Vegetation zone similarity, expressed as elevational distance and vegetation cover (NDVI residualized on elevation), and geographic proximity also showed positive associations with ASV sharing rate, though with smaller effect sizes than phylogenetic relatedness. The importance of phylogenetic relatedness as a predictor could also be partially explained by diet similarity among related species; dietary traits were found to be phylogenetically conserved among related host species (Kmulti = 0.671, p = 0.0003, 9,999 permutations, **supplementary figure S7**), and were therefore confounded with phylogenetic relatedness and excluded as an independent predictor in the model (**supplementary figure S8)**.

### Replicated signatures of host-microbial codiversification across the bacterial phylogeny in two mountain ranges

In contrast to strain sharing across host species, we next examined whether microbial lineages diversify in parallel within host lineages over evolutionary time, a pattern known as codiversification. We investigated patterns of codiversification between host and bacterial lineages by testing phylogenetic nodes across the distal quarter of the bacterial phylogeny. We found 109 nodes showing a strong signal of codiversification (Mantel’s rho > 0.75 and p-adj < 0.05) widely spread across the whole bacterial phylogeny (**Fig. 3A**). At the phylum level, Bacillota B (formerly Firmicutes) had the highest proportions of ASVs in strongly codiversifying nodes, followed by Actinomycetota, while Bacillota (also formerly Firmicutes, but a distinct subclade from Bacillota_B) had the highest number of ASVs with no members showing a strong codiversification signal (**Fig. 3B**). Codiversifying nodes represented a small fraction of the total relative abundance of bacteria present across host species (**Fig. 3C**). As examples of strong codiversification, we highlight a node corresponding to *Enterohabdus* sp., which showed the highest Mantel’s rho (**Fig. 3D-E**), and *Desulfonispora thiosulfatigenes*, the node with the greatest number of tips (**supplementary figure S9**). As examples of nodes lacking evidence for codiversification (Mantel’s rho < 0.25), we highlight *Enterococcus* sp. (**Fig. 3F-G**) and *Lactobacillus* sp. (**supplementary figure S10**), representing the single most abundant ASV across the dataset. These examples illustrate that the absence of codiversification can arise both from extensive sharing of the same ASVs among host species and from cases where multiple closely related ASVs are distributed inconsistently across hosts, resulting in weak correspondence between bacterial and host phylogenetic structure. A full list of codiversifying nodes is provided in **supplementary table S4**. To assess whether the codiversification patterns were reproducible across the two geographically isolated mountain ranges, we repeated Mantel tests separately within each mountain. Of the 109 strongly codiversifying nodes, 108 remained statistically significant (p < 0.05) in both mountains, and 70 retained a strong correlation (Mantel’s rho > 0.75) in both, demonstrating that codiversification signals were highly consistent across geographic replicates (**supplementary figure S11**).

**Figure 3.**
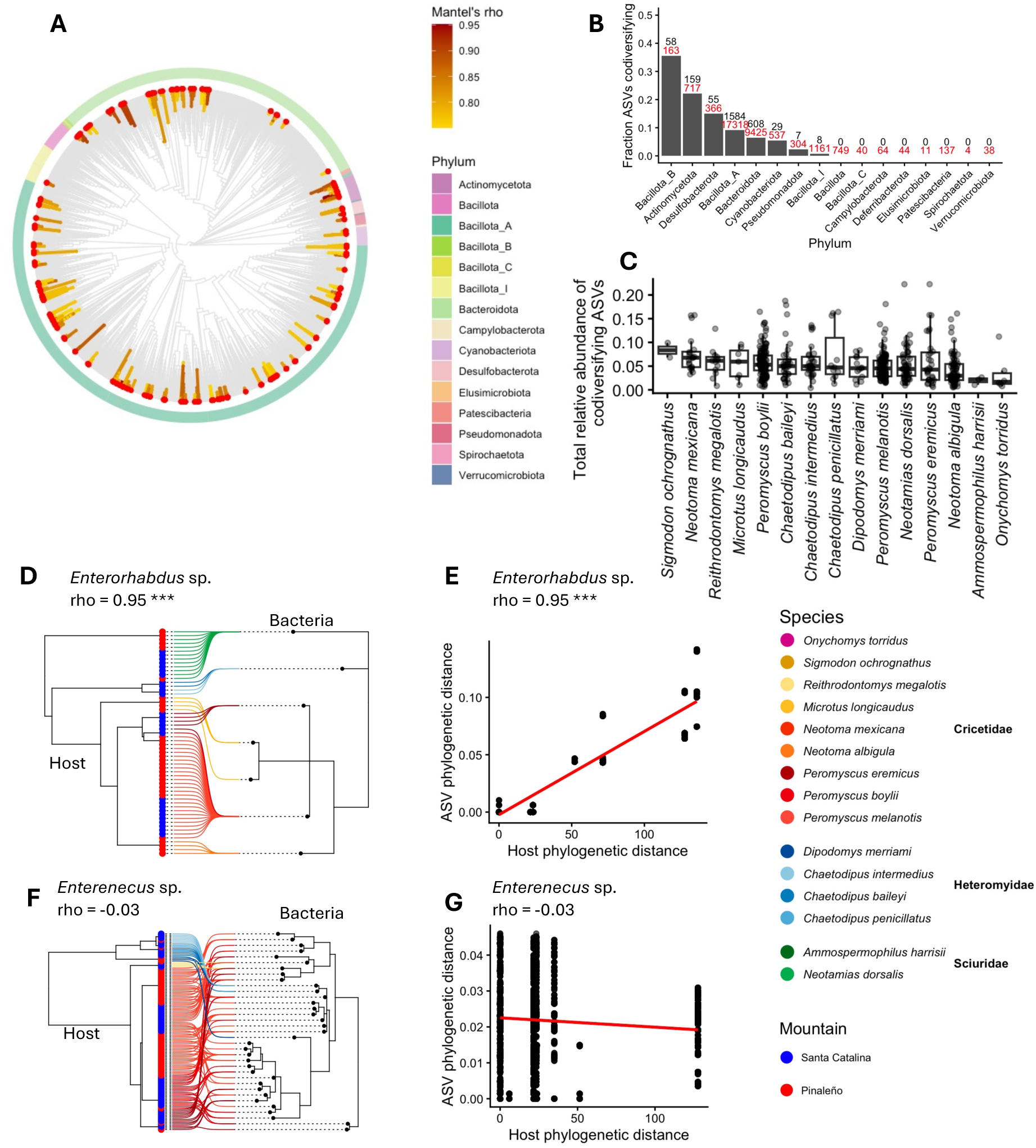
Strong codiversification signals are widespread across the bacterial phylogeny. (A) Circular cladogram of the bacterial phylogeny (10,000 randomly subsampled ASVs), with branches colored by Mantel’s rho indicating the strength of association between bacterial phylogeny and host-related factors. Tips are annotated by phylum (outer ring). (B) Bar plot showing the fraction of ASVs exhibiting significant codiversification signals within each bacterial phylum; numbers above bars indicate the total number of codiversifying ASVs (in black) and the total number of ASVs per phylum (in red). (C) Sum of the relative abundances of all ASVs showing strong codiversification per species, ordered by median value. (D and F) Tanglegram plots showing the relationship between host phylogeny (left) and bacterial phylogeny of selected nodes (right) for two bacterial taxa as examples. Lines connect host samples to associated bacterial ASVs and are colored by host species. The tips in the host phylogeny are colored by mountain. (E and G) Scatterplots showing the pairwise relationships between host and bacterial phylogenies for the corresponding nodes in panels D and F, respectively (Mantel test, *** P < 0.001).

These codiversifying lineages were also distinctly host-restricted in their distribution across the rodent community. The absence of codiversification signal can arise either from extensive sharing of identical ASVs among host species or from inconsistent distributions of closely related ASVs across hosts. We therefore examined how occupancy patterns differed between strongly codiversifying (rho > 0.75, adjusted p-value < 0.05) and non-codiversifying taxa (rho < 0.25). ASVs exhibiting strong codiversification signals tended to occur across fewer host species than ASVs lacking evidence of codiversification (**Fig. 4A**), indicating that codiversifying taxa were generally more host-specific. In contrast, the codiversification category was not significantly associated with sample-level occupancy (**Fig. 4B**), suggesting that codiversifying ASVs were not necessarily less prevalent across individuals. Despite occurring across fewer host species, strongly codiversifying ASVs exhibited higher average occupancy within host species (**Fig. 4C**). These patterns suggest that codiversifying taxa are specialists that achieve high prevalence within a narrow set of host lineages, a hallmark of host specialization, rather than generalists that persist broadly across the rodent community.

**Figure 4.**
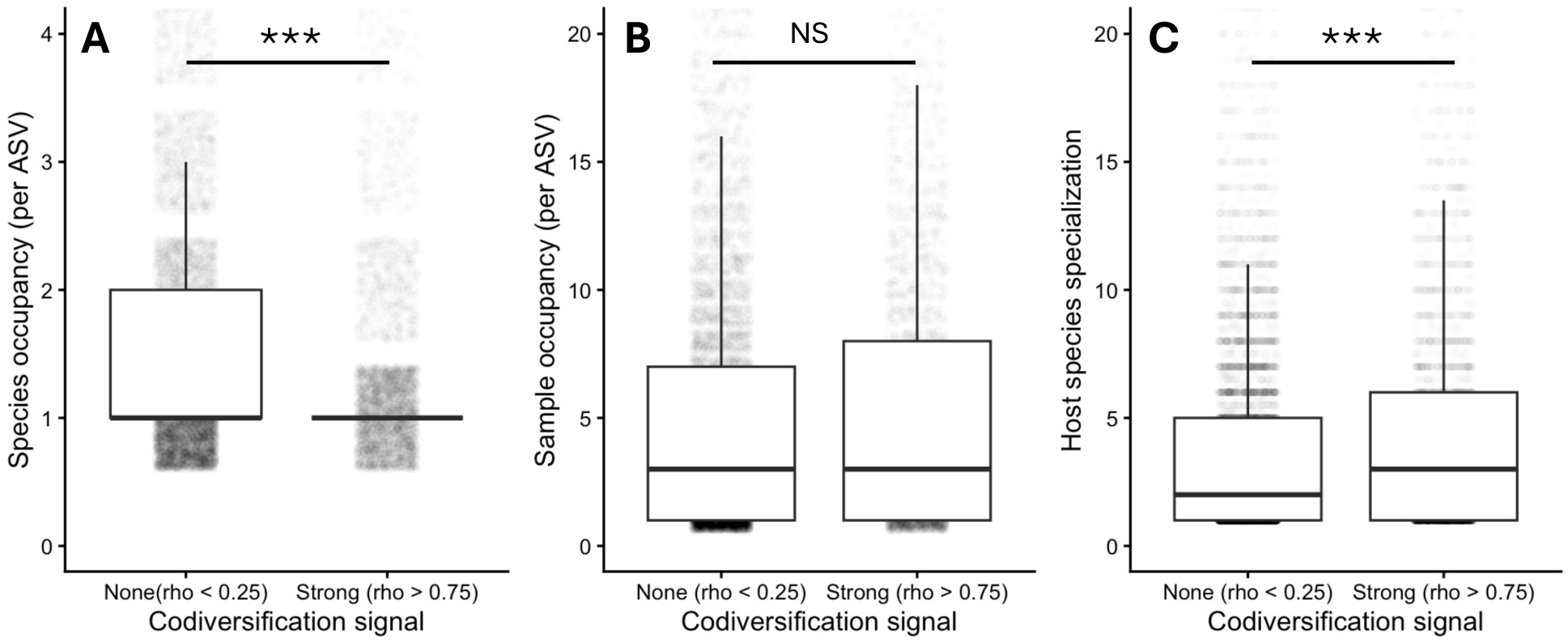
Associations between codiversification signal strength and metrics of occupancy (number of host species or samples colonized per ASV). (A) ASVs with no codiversification signal (rho < 0.25) exhibited significantly higher species occupancy than ASVs with a strong codiversification signal. (B) Sample occupancy did not differ significantly between codiversification categories. (C) ASVs with a strong codiversification signal showed significantly greater species specialization, measured as total sample occupancy normalized by species occupancy (P-values from the Wilcoxon rank-sum test, *** P < 0.001, NS = not significant).

### Microbial traits drive host specificity and interspecific strain sharing

To understand what drives codiversification and host specificity, we examined whether codiversifying lineages were enriched for traits reflecting adaptation to the gut environment and reduced capacity for environmental persistence. We grouped ASVs showing significant correlations by family and, for each family, calculated the fraction of ASVs falling within nodes exhibiting strong codiversification signals. Families represented by fewer than 10 ASVs across the dataset were excluded from the analysis. Within the remaining families, the proportion of ASVs associated with codiversifying nodes varied substantially (**Fig. 5A**). We then plotted these fractions against the predicted trait values for each family (**Fig. 5B-G**). To assess statistical significance, we compared the observed trait values of codiversifying taxa against null distributions generated by randomly assigning trait values across families (10,000 permutations). Codiversifying taxa showed evidence of adaptation to the host gut environment: obligate anaerobes were strongly over-represented among codiversifying taxa (99.3% vs. 81.7% null; p = 0.0002, **Fig. 5B**), and predicted optimal growth temperatures (OGT) clustered near host body temperature (mean deviation from 37°C of 1.47°C vs. 3.70°C under the null; p = 0.015, **Fig. 5E**). Contrary to expectations, sporulation was elevated among codiversifying lineages (27% versus 14% under the null; p = 0.03, **Fig. 5C**). Notably, codiversifying nodes were restricted to families with mean genome sizes between 1 and 3 Mb; however, permutation analyses showed that this pattern did not differ from expectations under the null model (p = 0.58; **Fig. 5D**). GC content likewise did not differ from the null (p = 0.64, **Fig. 5F**), and codiversifying taxa were more frequently Gram-positive (71% vs. 59% under the null), though this difference was not significant (p = 0.65, **Fig. 5G**). These traits themselves show strong phylogenetic clustering (anaerobic tolerance: D = −1.55, p < 0.001; sporulation: D = −1.95, p < 0.001; optimal growth temperature: λ = 0.69, p = 0.009), so we cannot rule out the possibility that shared ancestry, rather than codiversification itself, contributes to the trait associations reported above.

**Figure 5.**
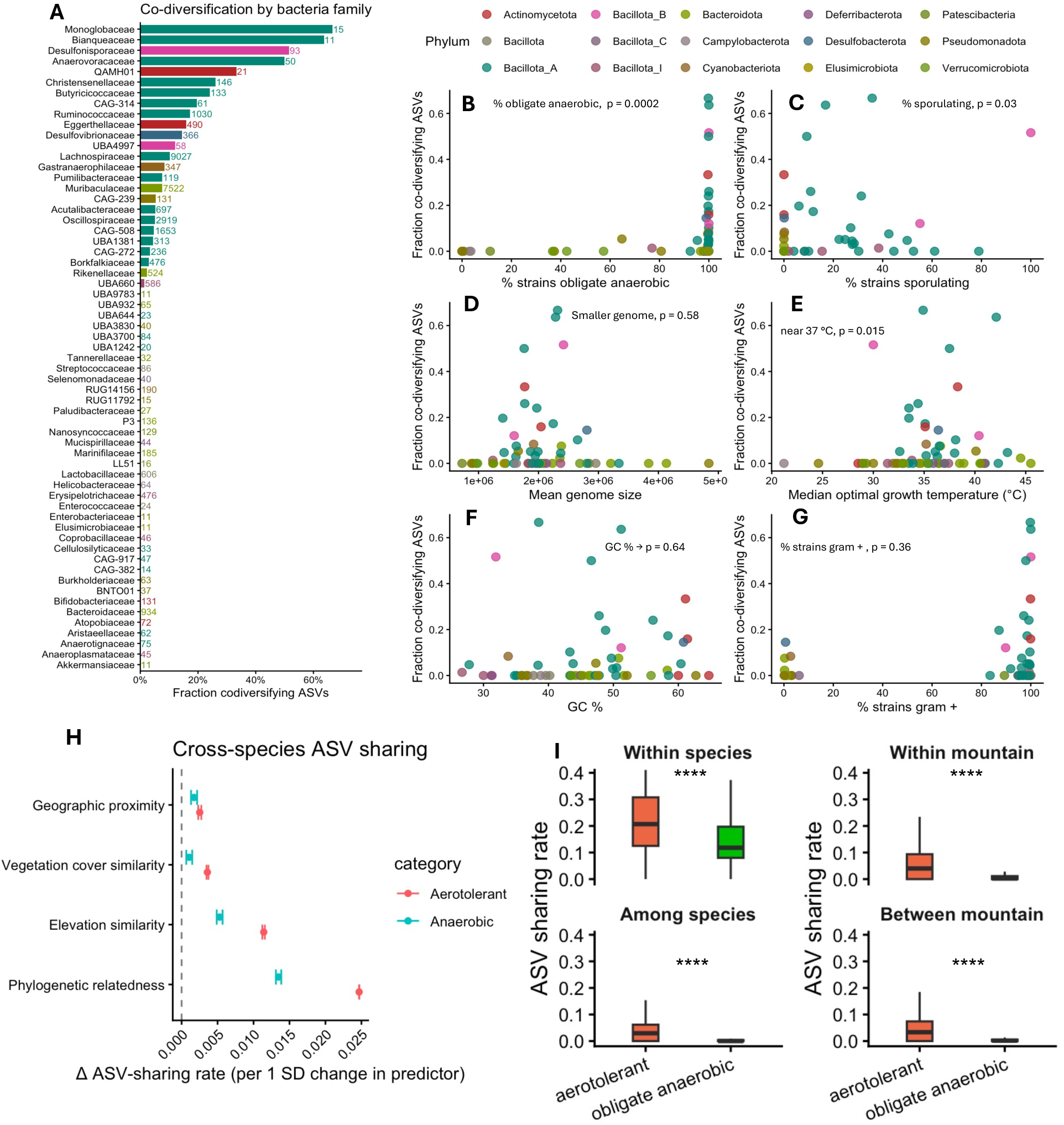
Taxonomic distribution and ecological correlates of codiversifying microbial ASVs. (A) Microbial families with at least 10 ASVs, with horizontal bars indicating the fraction of ASVs within each family identified as strongly codiversifying (rho > 0.75). Families are colored by phylum. The number of ASVs is shown next to the bars. (B-G) Relationships between the fraction of codiversifying ASVs and microbial ecological/genomic traits across families, including (B) the percentage of obligately anaerobic strains per family as reported in the MetaTrait DB, (C) the percentage of sporulating strains, (D) mean genome size, (E) median optimal growth temperature, (F) median GC %, and (G) the percentage of strains reported as Gram-positive. Each point represents a microbial family. P-values shown in each panel were derived from permutation tests (10,000 permutations) comparing the observed trait value against a null distribution generated by randomly reassigning traits across families; tests were one-sided where a directional hypothesis was specified (enrichment in obligate anaerobes, proximity to host body temperature of 37 °C, and a narrower genome-size range) and two-sided otherwise (GC content, sporulation, Gram-positive status). (H) Results of Bayesian multi-membership regression models estimating the effect of phylogenetic relatedness, elevational similarity, vegetation cover similarity (residuals of NDVI regressed on elevation) and geographic distance on pairwise ASV sharing rates across host species pairs, shown separately for aerotolerant bacteria and obligate anaerobes. Larger values indicate a stronger association between each predictor and the probability that two individuals share a given bacterial strain. Points represent posterior mean estimates; error bars indicate 95% credible intervals. (I) ASV sharing rates stratified by bacterial oxygen tolerance (aerotolerant vs. obligately anaerobic), faceted by host species identity (top) and mountain range co-occurrence (bottom). P-values were obtained by a permutation test comparing the observed difference in mean Sørensen similarity with a null distribution generated by randomly assigning aerotolerance labels among similarity values (9,999 permutations; **** P ≤ 0.0001).

We next examined whether the same microbial traits predict host occupancy, the number of host species in which an ASV was detected, as a continuous measure of interspecific strain sharing at the ASV rather than family level. We assigned traits related to environmental persistence and gut dependence, including aerotolerance, sporulation capacity, genome size, GC content, Gram stain, and OGT, to each ASV and modeled their association with host species occupancy using a generalized linear model (full model output in **supplementary table S5, supplementary figure S12**). Aerotolerant ASVs showed the strongest positive association with host species occupancy (β = 0.28, P < 0.001; detailed model results in Table S5). Genome size, OGT, and GC content all exhibited significant nonlinear relationships (quadratic terms, all P < 0.001), indicating that ASVs with more extreme trait values occupied more host species than those with intermediate values. Gram stain was also significantly associated with occupancy, although the magnitude of the association was small (β = 0.06, P < 0.001). The effect of sporulation was not significant (P = 0.77). Collectively, these results suggest that both environmental transmission potential and host-species-specific ecological filtering shape host occupancy patterns. However, when the analysis was repeated using a phylogenetic generalized linear mixed model that accounted for shared evolutionary history among the 5,000 most abundant ASVs, none of the fixed effects remained statistically significant (**supplementary table S6**), once again indicating that these trait associations are not independent of phylogenetic relatedness.

Finally, to confirm the role of aerotolerance in interspecific transmission patterns, we re-ran the Bayesian multilevel membership regression models separately for predicted aerotolerant and obligately anaerobic members of the community (**Fig. 5H**). Aerotolerant bacteria were shared at substantially higher rates than obligate anaerobes across all pairwise comparisons, including same- and different-species and same- and different-mountain pairs (**Fig. 5I**). Together, these three complementary analyses suggest that aerotolerance facilitated persistence outside the host and promoted colonization across host species, while obligate anaerobes with optimal growth temperatures near host body temperature (37°C) were more likely to diversify in concert with their host lineages.

## Discussion

A longstanding question in microbiome research is what determines whether a gut microbial lineage remains host-specific or is shared across multiple host species. We found that host phylogeny, rather than shared environment, is the strongest predictor of strain sharing, that signatures of host-microbial codiversification are widespread across the bacterial phylogeny and independently replicated across two mountain ranges, and that these patterns are associated with predicted microbial traits related to transmission potential and environmental persistence.

Species-specific microbiomes were strongly maintained even across substantial geographic separation of the two mountain ranges. Multiple mechanisms likely contribute to this pattern, including ecological selection by the host environment (Mazel et al., 2018), diet and behavioral similarity among related species (Youngblut et al., 2019), and stochastic transmission biased by host evolutionary history, consistent with the framework proposed by Kohl (2020) for phylosymbiosis. Social transmission may further reinforce species-specific microbiomes, particularly for bacterial taxa that have limited ability to survive outside host-associated environments (Debray et al., 2026; Petrullo et al., 2025; Raulo et al., 2024). Coprophagy, for example, is a known vector of microbial dispersal in rodents (Bo et al., 2020), and conspecifics likely have greater opportunities for this behavior given shared burrow and home range use. Critically, members of the same host species share a longer history of social contact and vertical and familial transmission than members of different species, and preferential within-species transmission of gut microbes over evolutionary time is therefore consistent with the host-specific strain distributions we observe.

Overall, microbial strain sharing across host species was limited, with only a small proportion of ASVs shared among species. Nevertheless, the extent of strain sharing varied considerably across species pairs, suggesting that some host species (*e.g.*, members of the Heteromyidae family) are more likely to exchange microbial lineages than others. To investigate the factors driving strain sharing, we modeled it as a function of phylogenetic relatedness, vegetation similarity, and geographic distance. When strain sharing was modeled explicitly, host phylogenetic relatedness emerged as the strongest predictor, consistent with host filtering as the primary driver of phylosymbiosis (Kohl 2020), and with previous studies identifying host phylogeny as the strongest determinant of strain sharing in wild rodents (Brown et al., 2023; Weinstein et al., 2021). Geographic proximity showed the weakest effect at the spatial scale of our study, which encompasses diverse rodent species in sympatry. In contrast, within species or closely related populations spanning broader geographic scales, geographic separation can become a primary driver: in continental-scale studies of deer mouse populations across the North American continent, isolation by distance was found to be the strongest determinant of strain divergence, exceeding the effect of phylogeny (Dillard et al., 2026), and major geographic barriers such as rivers have been shown to promote microbial strain divergence (Blumstein et al., 2026). We also detected a moderate effect of environmental similarity. That is, species that share a more similar habitat in terms of vegetation also share a higher proportion of bacterial strains in their microbiome. This likely reflects several non-mutually exclusive mechanisms: overlapping diets among species feeding on similar vegetation (Weinstein et al., 2021; Youngblut et al., 2019), shared exposure to environmentally acquired microbes (H. He et al., 2026; Xia et al., 2025), and increased opportunities for interspecific contact within the same habitats (Perofsky et al., 2019).

While interspecific strain sharing was limited overall, signals of host-microbial codiversification were widespread across the bacterial phylogeny, consistent with a previous study in rodents (Sprockett et al., 2025), and strikingly, independently replicated across the two mountain ranges. Of the 109 strongly codiversifying nodes, 108 remained significant in both mountains independently, demonstrating a robustness of signal that is rarely achievable in field-based microbiome studies. This replication across populations diverging for 10,000 - 50,000 years strengthens the inference that codiversification reflects parallel host and microbial divergence rather than convergent responses to similar host environments. At the family level, the codiversifying lineages were dominated by obligate anaerobic fermenters, including Lachnospiraceae (Vacca et al., 2020), Ruminococcaceae (Biddle et al., 2013), Oscillospiraceae (Gophna et al., 2017), Butyricicoccaceae (Boesmans et al., 2018) and Monoglobaceae (Yüksel et al., 2025), which are predominantly short-chain fatty acid (SCFA) producers, as well as taxa involved in sulfur and bile-acid metabolism such as Desulfovibrionaceae, Desulfonisporaceae and Eggerthellaceae (Denger et al., 1999; Ridlon et al., 2006; Song et al., 2019). Christensenellaceae also showed strong codiversification signals; this family is among the most heritable bacterial families known from human studies (Goodrich et al., 2017; Waters & Ley, 2019). Codiversifying taxa were also distinctly host-specific, occurring across fewer host species but achieving higher prevalence within the host lineages they colonized, a pattern indicative of host specialization rather than generalist persistence.

The convergence of codiversification patterns between our rodent system and the human gut microbiome suggests a conserved mechanism of microbial dispersal limitation across mammalian hosts. In humans, codiversifying gut bacteria tend to be oxygen intolerant and temperature sensitive, with reduced genome sizes reflecting host dependency (Suzuki et al., 2022). We observed strikingly similar patterns in our rodent communities: codiversifying taxa were strongly enriched for obligate anaerobes with predicted optimal growth temperatures near host body temperature, suggesting poor survival outside the host environment. Genome reduction in codiversifying taxa did not reach statistical significance, though this may partly reflect the inclusion of intracellular bacterial parasites such as Nanosyncoccaeae (W. He et al., 2024) and UBA1242 (Begmatov et al., 2025), whose extremely small genomes could obscure the pattern. The bacterial families dominating codiversification in our dataset, particularly Lachnospiraceae and Ruminococcaceae, overlap with codiversifying taxa identified in humans, including butyrate producers (Suzuki et al., 2022). Whether enrichment of SCFA-fermenting lineages among codiversifying taxa is a general feature of mammalian host-microbial codiversification remains to be tested, but the convergence across rodents and humans is consistent with this possibility. In contrast, aerotolerant taxa showed substantially higher interspecific strain sharing rates across all pairwise comparisons, suggesting they are better equipped to persist outside the host and colonize new species. This is consistent with other rodent systems, where aerotolerant taxa are more frequently transmitted through indirect and social routes within populations (Petrullo et al., 2025; Raulo et al., 2024). Together, microbial traits reflecting environmental persistence and gut adaptation predicted both codiversification status and interspecific strain sharing rate, suggesting that the same physiological axes governing within-population transmission dynamics also shape evolutionary outcomes across deeper timescales.

Our study has limitations worth considering. First, we did not collect dietary data, and because closely related host species often share similar diets, some of the observed associations between phylogenetic relatedness and microbial strain sharing may partly reflect shared feeding ecology. However, since diet preference itself is partly a product of host evolutionary history, phylogenetic relatedness as a predictor already captures a component of dietary similarity, and future studies integrating dietary data will help partition these contributions more precisely. Our analyses relied on single-marker-gene sequencing, specifically full-length 16S rRNA amplicons of approximately 1,500 bp, which provided substantially greater strain-level resolution than short-read approaches (Johnson et al., 2019), and were sufficient to detect robust codiversification signals within bacterial species. However, single-marker-gene phylogenies may not always reflect whole-genome evolutionary histories (Hassler et al., 2022), and microbial traits were also predicted from single marker gene taxonomy rather than from genomes or functional assays. Furthermore, these traits are phylogenetically conserved, making it difficult to fully distinguish whether they independently drive codiversification and transmission patterns or instead characterize bacterial clades inherently more prone to host restriction of dispersal. The patterns reported here should therefore be interpreted as broad evolutionary associations rather than definitive evidence of parallel host and microbial diversification. Future studies should also expand beyond traits related to environmental persistence to examine microbial metabolic pathways, which may determine how microbial lineages establish and maintain long-term associations with particular hosts. The full-length 16S data nonetheless provide a powerful starting point for identifying candidate microbial taxa and samples for targeted follow-up using direct shotgun metagenomics, probe-based target enrichment, or culture-based whole-genome sequencing, all of which would enable high-resolution phylogenetic and functional analyses at a lower cost than applying untargeted metagenomics across all samples. Such data, combined with host genomic data, would also allow direct tests of whether bacterial lineages that codiversified across host species continue to track host population structure within species by assessing concordance between host and microbial population genetic structure.

In summary, host evolutionary history was the primary determinant of gut microbial strain sharing across sympatric Sky Island rodent communities, with signals of codiversification widespread across the bacterial phylogeny and independently replicated across two geographically isolated mountain ranges. Strikingly, the same mechanistic signature emerged across systems: microbial traits reflecting reduced environmental persistence and gut adaptation predicted codiversification in rodents and have been associated with codiversification within human populations (Suzuki et al. 2022). Together, these findings suggest that physiological adaptation to the gut, including limited aerotolerance and optimal growth near host body temperature, represents a conserved axis of microbial dispersal limitation shaping the evolution of mammalian gut symbioses. Investigating how barriers to microbial transmission generate host specificity and, over evolutionary time, codiversification offers a framework for understanding how microbial communities respond to environmental change, host population fragmentation, and emerging disease dynamics.

## Data availability

Raw sequencing data are available in the NCBI Sequence Read Archive under the bioproject ID PRJNA1514934. Code is available at https://github.com/gabridinosauro/sky-islands-analysis

## Acknowledgements

We thank members of the Upham lab, Pasch lab, and Suzuki lab (especially Danielle Blumstein, Emerson Fongenyam, and Thomas Archer-Black) who helped collect the samples, and members of the ASU Biodesign Center for Health Through Microbiomes for helpful comments and support during the development of this project. The project and T.A.S. were supported by the National Institutes of Health (R35 GM160076). N.S.U. was supported by the National Institutes of Health (1R21AI164268, 1R35GM156919).

