## Supplementary Figure for "Conserved patterns of mammal–microbe codiversification and interspecific bacterial sharing"

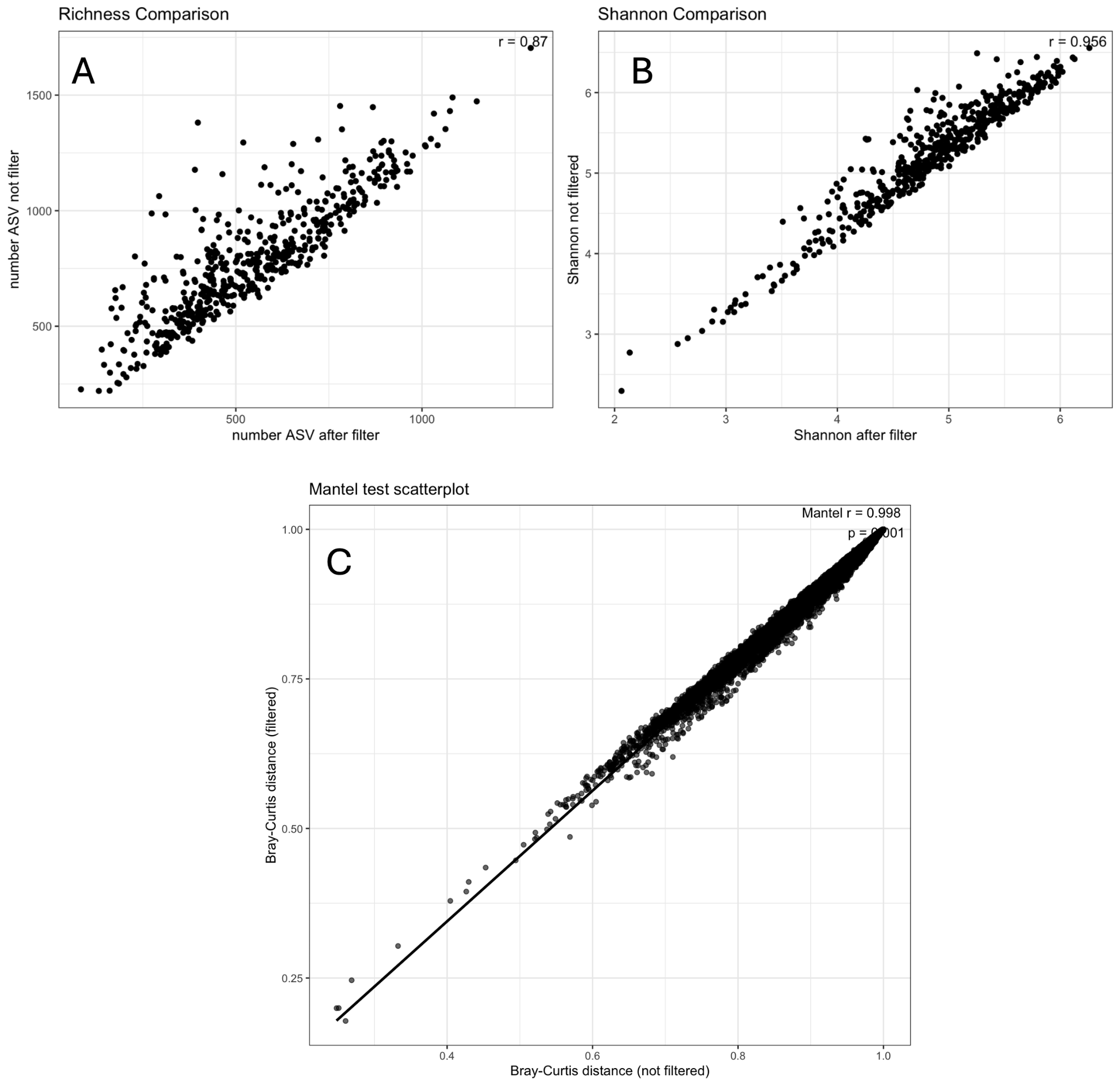

**Supplementary Figure S1.** Comparison between alpha and beta diversity patterns using the full ASV table and the filtered ASV table, in which ASVs were retained if they had at least 50 total counts across all samples and reached a minimum abundance of at least 10 reads in at least one sample. (A) Comparison of observed richness (number of ASVs) calculated from the filtered and unfiltered datasets. (B) Comparison of Shannon diversity values calculated from filtered and unfiltered datasets. (C) Comparison of Bray–Curtis dissimilarity matrices derived from the filtered and unfiltered datasets. Results demonstrate that filtering had minimal impact on the inferred beta-diversity relationships among samples.

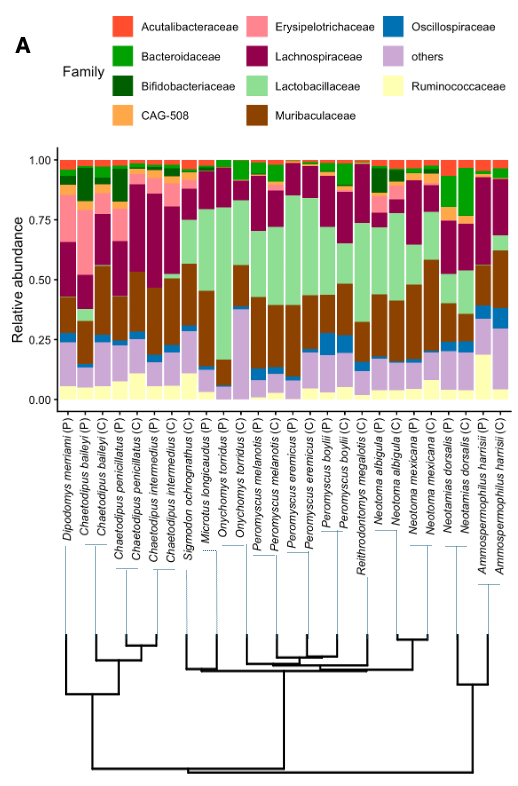

**Supplementary Figure S2.** Relative abundance of bacterial families across host species, with samples grouped by species and site (P = Pinaleño Mountains; C = Santa Catalina Mountains). Colors indicate the dominant microbial families (the top 10 across the whole dataset), with less abundant groups grouped into “others”. The phylogenetic tree below shows evolutionary relationships among host species, allowing comparison between host relatedness and microbiome composition.

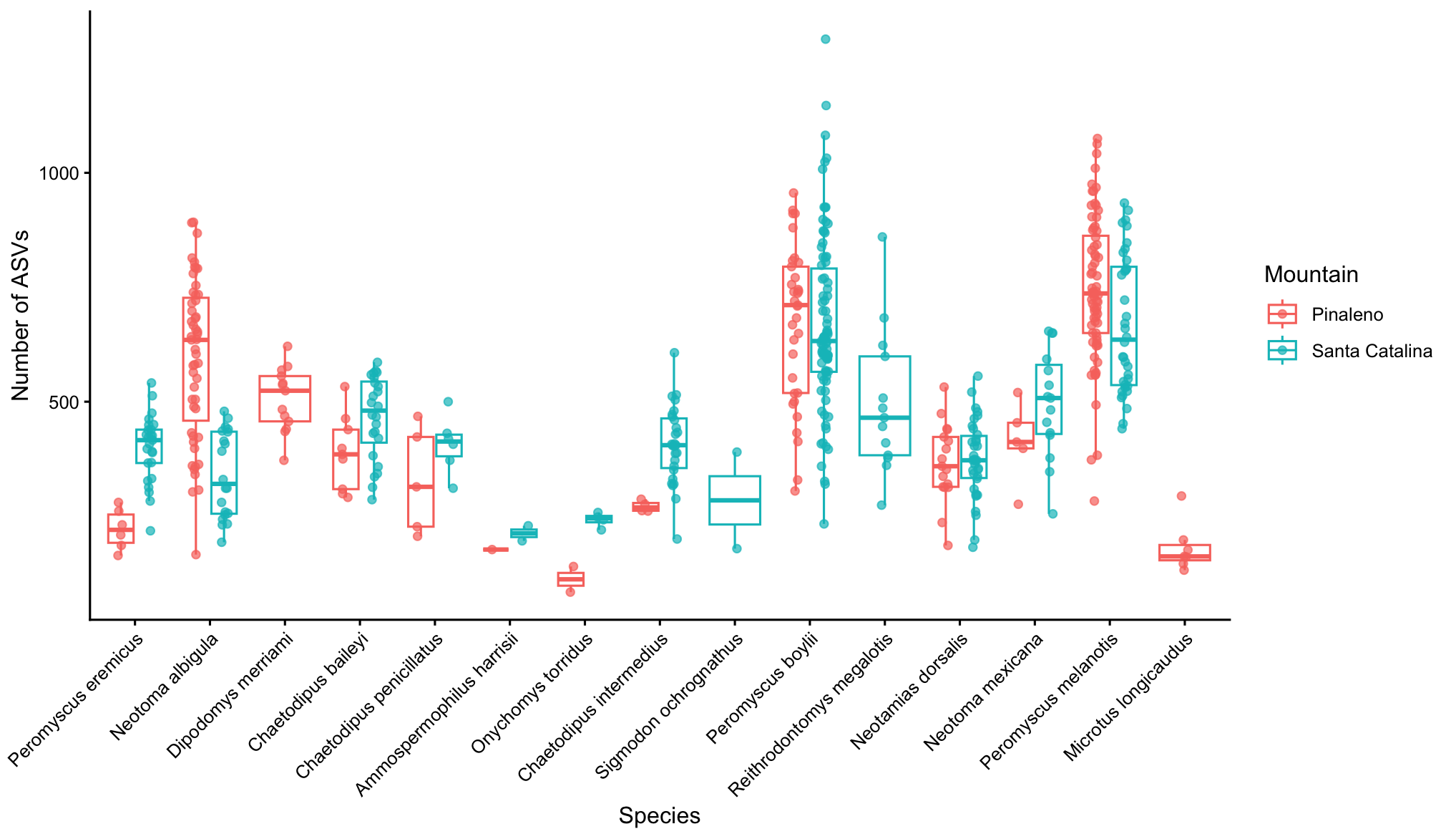

**Supplementary Figure S3.** Alpha-diversity across host species and mountain ranges. Boxplots show the distribution of ASV richness per individual for each of the 15 rodent species sampled, grouped by mountain range (Pinaleno in red and Santa Catalina in blue), with species ordered by their mean elevation of occurrence.

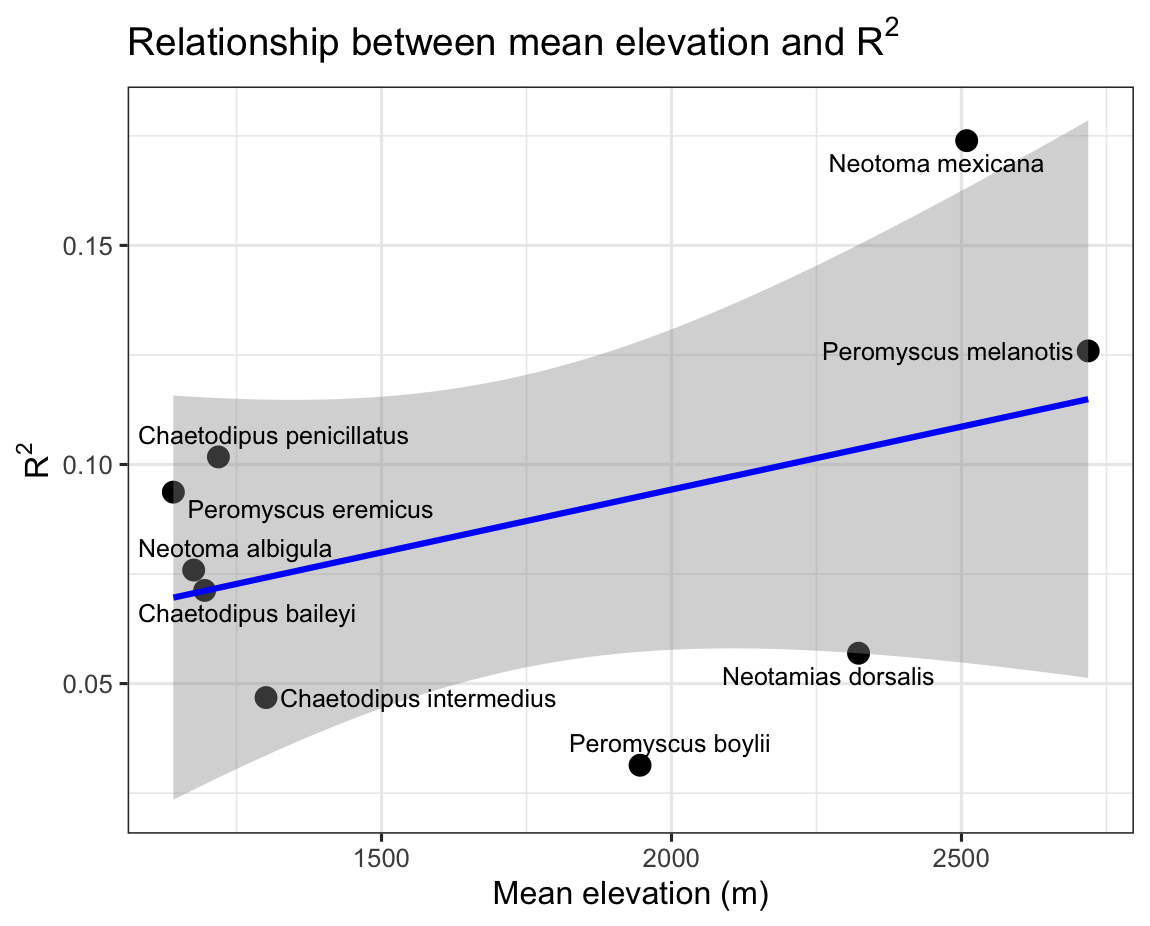

**Supplementary Figure S4.** Correlation between host species mean elevation and PERMANOVA R2 ( r = 0.45, p = 0.25).

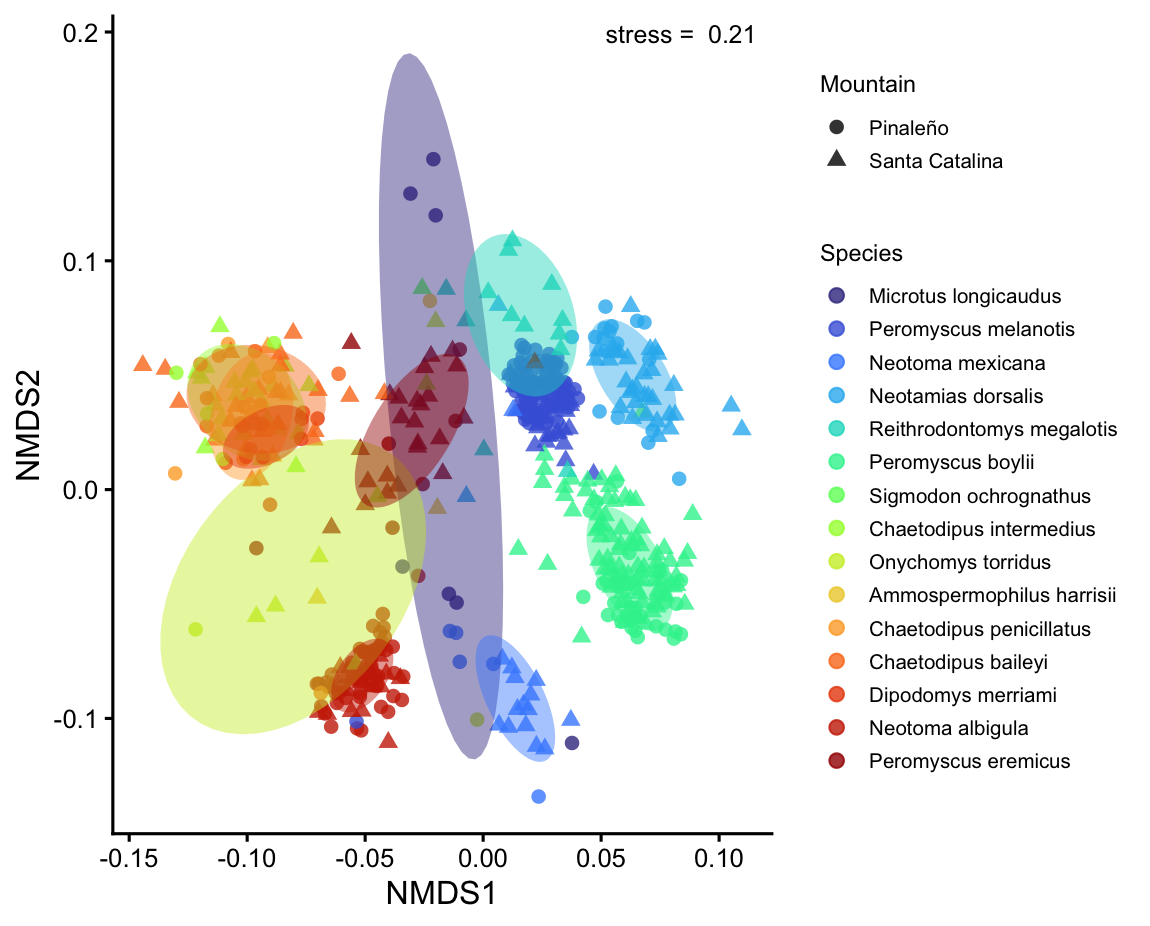

**Supplementary Figure S5.** Permutational multivariate analysis of variance (PERMANOVA) based on Bray–Curtis dissimilarity. Host species explained the largest proportion of variation (R² = 0.134, *F* = 5.94, *P* = 0.001), followed by mountain of origin (R² = 0.008, *F* = 5.11, *P* = 0.001), residual NDVI (R² = 0.006, *F* = 3.57, *P* = 0.001), and elevation (R² = 0.004, *F* = 2.36, *P* = 0.001), all of which were statistically significant. Colors indicate host species ordered by mean elevation of occurrence and symbols denote mountain range.

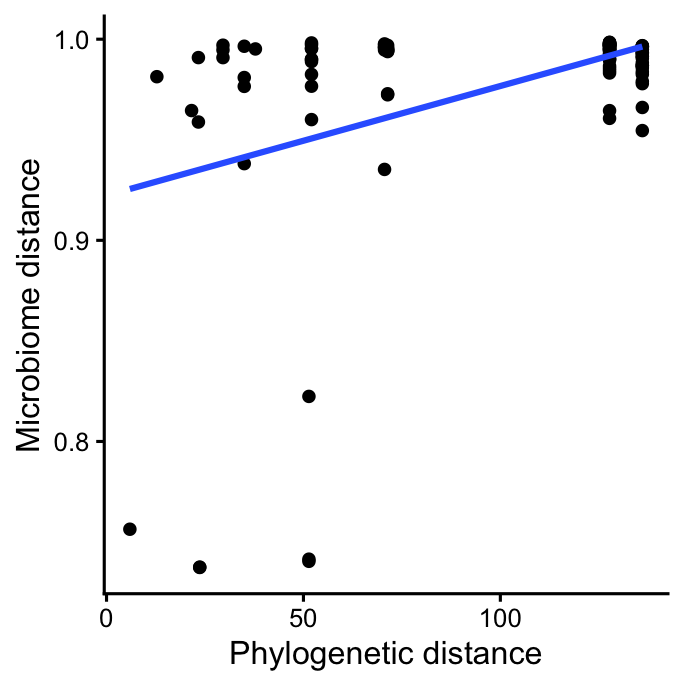

**Supplementary Figure S6.** Relationship between host phylogenetic distance and gut microbiome Sørensen–Dice dissimilarity among host species. For each host species, the average microbiome composition was calculated by pooling all individuals and generating an average Sørensen–Dice dissimilarity matrix across 100 rarefaction replicates (rarefaction depth: 10,000 reads). The blue line shows the fitted linear regression. A Mantel test confirmed a statistically significant positive correlation between host phylogenetic distance and microbiome Sørensen–Dice dissimilarity (Mantel rho = 0.198, *P* = 0.046, 9,999 permutations), indicating that more distantly related host species tend to harbor more dissimilar gut microbiomes, evidence of phylosymbiosis.

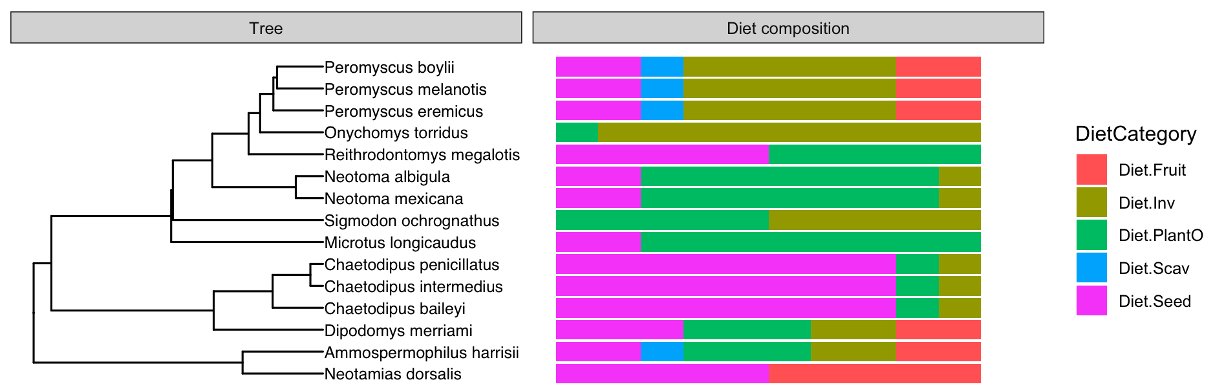

**Supplementary Figure S7.** Dietary composition and how it is related to phylogenetic distance among sky island rodent species. Left side, phylogenetic tree of the study species, on the right stacked bar plots of proportional diet categories (Elton Traits: seed, plant/other, invertebrate, fruit, scavenge) for each species. Diet composition appears conserved within genera (*Peromyscus*, *Chaetodipus*, *Neotoma*) but varies more independently among more distantly related taxa.

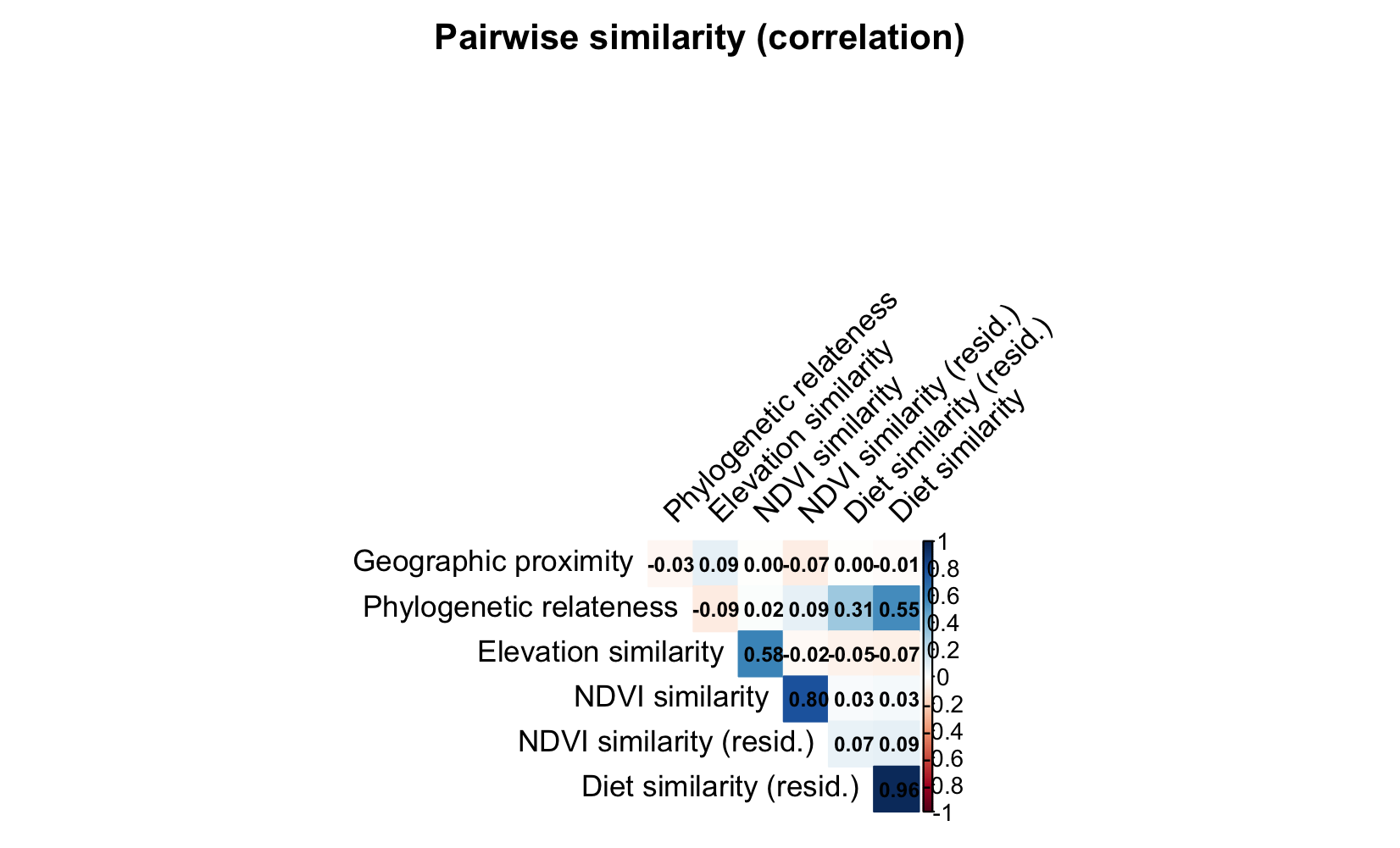

**Supplementary Figure S8.** Pairwise Pearson correlations among geographic, phylogenetic, elevation, NDVI, and diet distance metrics among paired interspecific comparisons. Color and value indicate Pearson's *r* (blue = positive, red = negative).

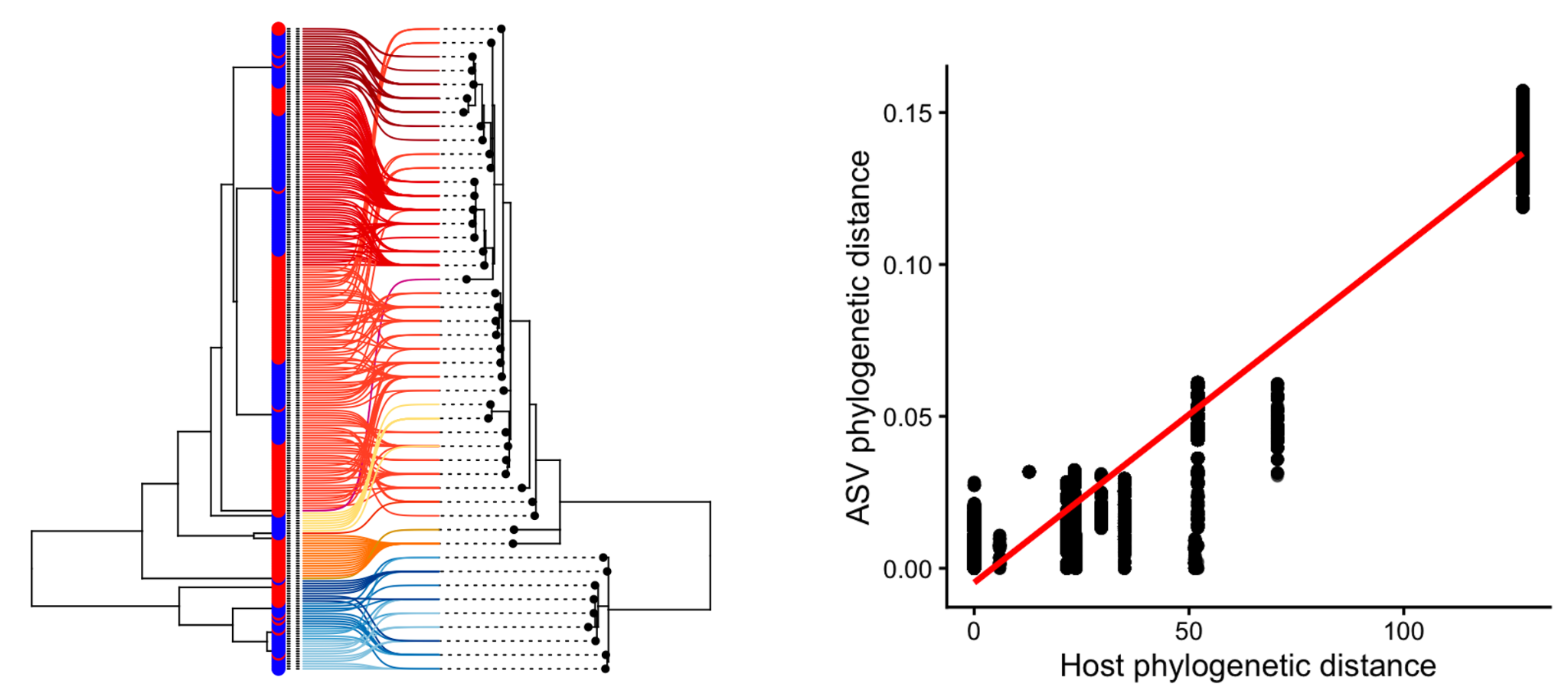

**Supplementary Figure S9.** Codiversification analysis of *Desulfonispora thiosulfatigenes* clade as an example of a strong codiversification signal. The left panel shows a tanglegram displaying the relationship between host phylogeny (left tree) and bacterial phylogeny (right tree) of the selected nodes. Lines connect host samples to their associated bacterial ASVs and are colored by host species (same color codes as Figure 3). Tip colors on the host phylogeny indicate mountain range (blue: Santa Catalina, red: Pinaleño). The right panel shows a scatterplot of pairwise host and bacterial phylogenetic distances for the visualized nodes (Mantel rho = 0.90, P < 0.001).

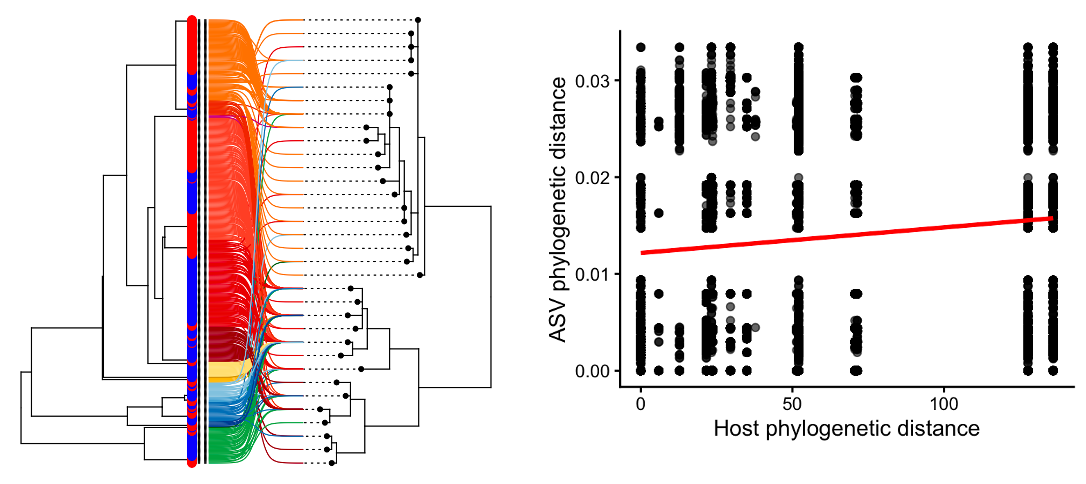

**Supplementary Figure S10.** Codiversification analysis of *Lactobacillus* sp. clade as an example of a significant but weak codiversification signal. The left panel shows a tanglegram displaying the relationship between host phylogeny (left tree) and bacterial phylogeny (right tree) of the selected nodes. Lines connect host samples to their associated bacterial ASVs and are colored by host species (same color codes as Figure 3). Tip colors on the host phylogeny indicate mountain range (blue: Santa Catalina, red: Pinaleño). The right panel shows a scatterplot ofpairwise host and bacterial phylogenetic distances for the visualized nodes (Mantel rho = 0.24, P < 0.01).

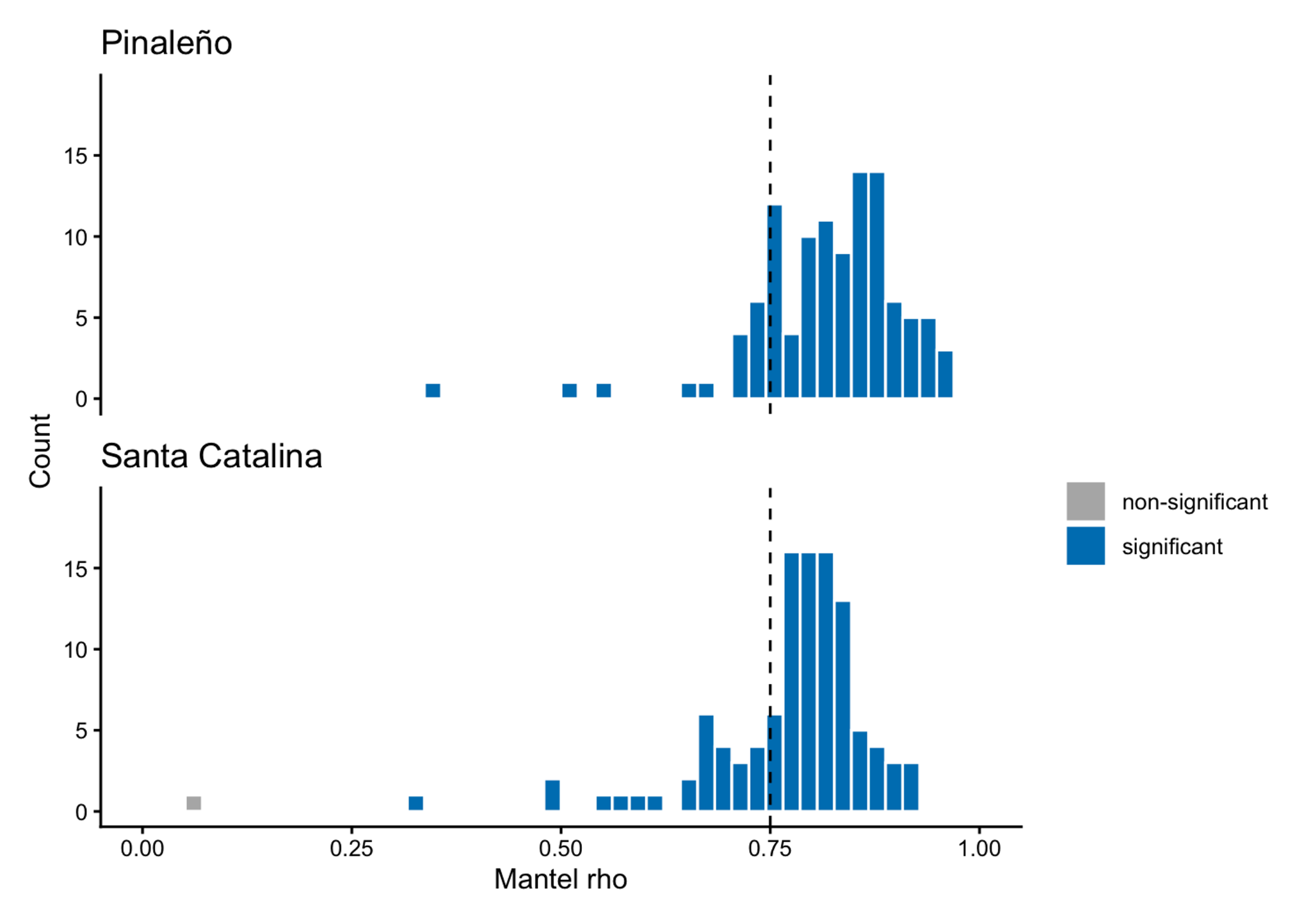

**Supplementary Figure S11.** Replicated host-microbial codiversification signals across independent mountain ranges. Histograms show the distribution of Mantel correlation coefficients (Mantel’s rho) between host and bacterial phylogenetic distances for the 109 strongly codiversifying nodes (Mantel rho> 0.75, adjusted P < 0.05), analyzed separately for for Pinaleño (top) and Santa Catalina (bottom) mountains. Bar colors indicate significant (blue; permutation *p* < 0.05) and non-significant (gray) results. Of the 109 nodes, 108 remained statistically significant in both mountain ranges independently, indicating that the host-microbial codiversification signals are robustly replicated across geographically isolated populations.

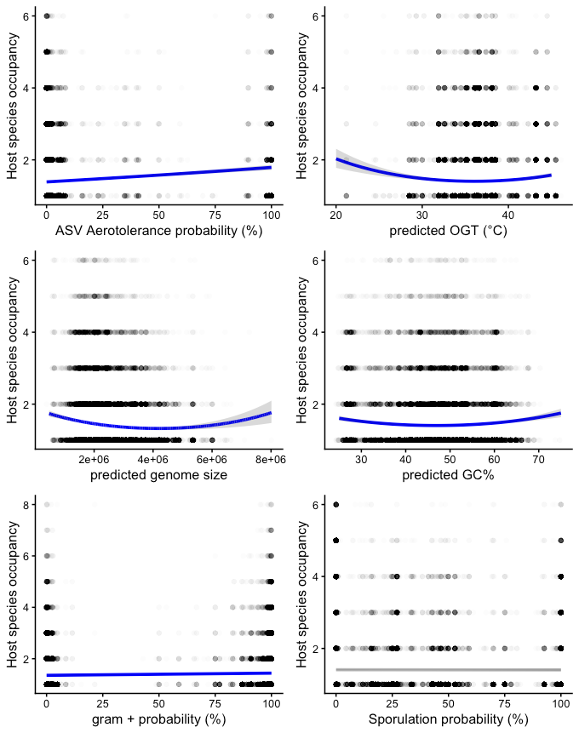

**Supplementary Figure S12.** Associations between microbial traits and host species occupancy (the number of host species in which an ASV was detected). Trend lines show fitted relationships from generalized linear models, with shaded areas representing 95% confidence intervals. Aerotolerance showed the strongest positive association with occupancy. Genome size, optimal growth temperature (OGT), and GC content exhibited significant nonlinear relationships, with ASVs at more extreme trait values occupying fewer host species than those with intermediate values. The probability of being Gram-positive showed a weak positive association with occupancy, while sporulation showed no significant association. See Table S5 for test statistics and p-values.
