## Supplementary table for "Conserved patterns of mammal–microbe codiversification and interspecific bacterial sharing"

**Supplementary table S1** (see excel file)

**Supplementary table S2.** Results of PERMANOVA analyses testing for genetic differentiation among mountain ranges within each species. Shown are sample size (n), number of mountain ranges included in the analysis, PERMANOVA R², pseudo-F statistic, and permutation-based p-value for each species.

| **Species** | **n** | **Mountains** | **R²** | **F** | **p** |
| --- | --- | --- | --- | --- | --- |
| *Peromyscus eremicus* | 31 | 2 | 0.094 | 2.998 | 0.001 |
| *Neotoma albigula* | 71 | 2 | 0.076 | 5.668 | 0.001 |
| *Chaetodipus baileyi* | 33 | 2 | 0.071 | 2.378 | 0.001 |
| *Chaetodipus penicillatus* | 11 | 2 | 0.102 | 1.019 | 0.371 |
| *Chaetodipus intermedius* | 31 | 2 | 0.047 | 1.424 | 0.004 |
| *Peromyscus boylii* | 119 | 2 | 0.031 | 3.789 | 0.001 |
| *Neotamias dorsalis* | 51 | 2 | 0.057 | 2.958 | 0.001 |
| *Neotoma mexicana* | 20 | 2 | 0.174 | 3.789 | 0.001 |
| *Peromyscus melanotis* | 108 | 2 | 0.126 | 15.270 | 0.001 |

**Supplementary Table S3.** Table of Average marginal effects estimated from the Bayesian ordinal beta regression model predicting Sørensen similarity. Estimates are reported on the response scale and represent the average change in predicted Sørensen similarity associated with a two-unit of standard deviation increase in each predictor, with other variables held constant. Values are posterior means with 95% credible intervals.

| Predictor | Average marginal effect | 95% Credible Interval |
| --- | --- | --- |
| Phylogenetic relatedness (106 MI) | 0.01639 | 0.01626–0.01652 |
| Elevation similarity (1246 m) | 0.00631 | 0.00622–0.00642 |
| NDVI residuals (0.23) | 0.00171 | 0.00162–0.00181 |
| Geographic proximity (81 km) | 0.00169 | 0.00161–0.00176 |

**Supplementary Table S4** (see excel file)

**Supplementary Table S5.** Binomial generalized linear model predicting host species occupancy (the number of host species in which an ASV was detected) as a function of microbial traits. Continuous predictors (genome size, temperature, GC content) were z-score standardized prior to modeling; quadratic terms were included to account for nonlinear relationships with occupancy. Binary traits including aerotolerance, sporulation, and Gram stain were scaled on a 0–1 proportion scale representing the fraction of strains within each taxonomic group displaying that trait.

| Predictor | Estimate | SE | z | P-value |
| --- | --- | --- | --- | --- |
| Intercept | -2.32 | 0.008 | -287.4 | <0.001 |
| Aerotolerance | 0.28 | 0.023 | 12.06 | <0.001 |
| Sporulation | 0.004 | 0.014 | -0.28 | 0.77 |
| Genome size | -0.06 | 0.007 | -9.209 | <0.001 |
| Genome size² | 0.014 | 0.002 | 5.4 | <0.001 |
| Temperature | 0.007 | 0.004 | 1.65 | 0.1 |
| Temperature² | 0.011 | 0.002 | 6.39 | <0.001 |
| GC content | 0.004 | 0.004 | 0.84 | 0.4 |
| GC content² | 0.02 | 0.003 | 6.637 | <0.001 |
| Gram stain (gram-positive) | 0.06 | 0.011 | 5.89 | <0.001 |
| Statistic | Value |  |  |  |
| Null deviance | 25096 |  |  |  |
| Residual deviance | 24086 |  |  |  |
| AIC | 125183 |  |  |  |
| Observations | 46,566 |  |  |  |
| Missing observations removed | 1,216 |  |  |  |

**Supplementary table S6.** Results of the phylogenetic generalized linear mixed model (PGLMM) examining predictors of species occurrence. Continuous predictors (genome size, temperature, GC content) were standardized (z-scores), quadratic terms were included to test for potential nonlinear relationships between microbial traits and host species occupancy. Binary traits including aerotolerance, sporulation, and Gram stain were scaled on a 0–1 proportion scale representing the fraction of strains within each taxonomic group displaying that trait.

| Predictor | Estimate (β) | Std. Error | z | P-value |
| --- | --- | --- | --- | --- |
| Intercept | -2.442 | 0.177 | -13.791 | <0.001 *** |
| Aerobic | 0.303 | 0.191 | 1.583 | 0.113 |
| Sporulation | -0.035 | 0.062 | -0.568 | 0.570 |
| Genome size | -0.027 | 0.034 | -0.796 | 0.426 |
| Genome size² | 0.013 | 0.013 | 1.004 | 0.315 |
| Temperature | -0.012 | 0.032 | -0.372 | 0.710 |
| Temperature² | -0.0002 | 0.009 | -0.023 | 0.981 |
| GC content | -0.050 | 0.031 | -1.630 | 0.103 |
| GC content² | -0.013 | 0.020 | -0.671 | 0.503 |
| Gram-positive | 0.193 | 0.155 | 1.246 | 0.213 |
